# Meta-learning local synaptic plasticity for continual familiarity detection

**DOI:** 10.1101/2021.03.21.436287

**Authors:** Danil Tyulmankov, Guangyu Robert Yang, LF Abbott

## Abstract

Over the course of a lifetime, a continual stream of information is encoded and retrieved from memory. To explore the synaptic mechanisms that enable this ongoing process, we consider a continual familiarity detection task in which a subject must report whether an image has been previously encountered. We design a class of feedforward neural network models endowed with biologically plausible synaptic plasticity dynamics, the parameters of which are meta-learned to optimize familiarity detection over long delay intervals. After training, we find that anti-Hebbian plasticity leads to better performance than Hebbian and replicates experimental results from the inferotemporal cortex, including repetition suppression. Unlike previous models, this network both operates continuously without requiring any synaptic resets and generalizes to intervals it has not been trained on. We demonstrate this not only for uncorrelated random stimuli but also for images of real-world objects. Our work suggests a biologically plausible mechanism for continual learning, and demonstrates an effective application of machine learning for neuroscience discovery.

## Introduction

Every day, a continual stream of sensory information and internal cognitive processing causes lasting synaptic changes in our brains that alter our responses to future stimuli. It remains a mystery how neural activity and local synaptic updates coordinate to support distributed storage and readout of information and, in particular, how ongoing synaptic changes due to either new memories or homeostatic mechanisms do not interfere with previously stored information.

Memory research in theoretical neuroscience and machine learning has addressed these questions through modeling studies, but important features remain to be clarified. First, memories of an individual’s history are encoded in a one-shot manner – this is different from typical neural network models which use a prolonged incremental training process to learn a complex task. Such training algorithms use a global error signal and perform per-synapse credit assignment through knowledge of the entire network (Rumelhart et al., 1986), whereas biological synapses typically only have access to local pre- and postsynaptic activity (Hebb, 1949) and various modulatory signals (Frémaux and Gerstner, 2016; Gerstner et al., 2018). Second, biological synapses change continually in response to ongoing activity, whereas models commonly assume that synapses are fixed after training ends, such as the classical Hopfield network (Hopfield, 1982) and most deep neural networks (LeCun et al., 2015). Unregulated continual updating of synapses can cause catastrophic forgetting in which a network either erases previous memories (Kirkpatrick et al., 2017; Zenke et al., 2017) or renders stored information unreadable (Parisi, 1986). Recurrent neural networks are commonly used to perform tasks that involve memories sustained by neural activity (Elman, 1991; Hochreiter and Schmidhuber, 1997; Mante et al., 2013), however most memories are likely stored through synaptic potentiation and depression (Abbott and Nelson, 2000). Synaptic memory has sometimes been studied through an ideal observer approach (Benna and Fusi, 2016; Fusi et al., 2005) in which synaptic weights are directly accessible for readout, but biological organisms must read out synaptic storage through neuronal activations. In fact, the readout may not be a dedicated circuit as in attractor network models of memory (Hopfield, 1982), but rather manifested as a change in ongoing neural processing (Hasson et al., 2015). Finally, machine learning research often eschews biological plausibility and mechanistic understanding in favor of performance on benchmark tasks (Ba et al., 2016; Graves et al., 2014, 2016; Miconi et al., 2018, 2019).

Familiarity detection – identifying whether a stimulus has been previously encountered – is a simple and ubiquitous form of memory that serves as a useful testbed for addressing these issues. Classical studies have demonstrated that human recognition memory capacity for images is “almost limitless” in a two-alternative-forced-choice task with separate encoding and testing phases: retention follows a power law as a function of the number of items viewed (Standing, 1973). Pioneering theoretical work has shown that the number of memories stored by a familiarity detection network depends on the synaptic plasticity rule and, in the case of uncorrelated inputs, capacity can scale proportionally to the number of synapses (Bogacz and Brown, 2003). More recent behavioral work further demonstrates an impressive capacity in a continual setting, the error rate as a function of the number of intervening items exhibiting a “power law of forgetting” (Brady et al., 2008). Theoretical studies have shown that power-law forgetting is achievable by synapses with metaplasticity, using both uncorrelated inputs (Fusi et al., 2005) and face images (Ji-An et al., 2019). Neural signals of visual familiarity have been observed as reductions in responses to repeated presentations of a stimulus, a phenomenon known as repetition suppression (Grill-Spector et al., 2006; Meyer and Rust, 2018; Miller et al., 1991; Xiang and Brown, 1998). At the timescales relevant for this task – one-shot memorization on the order of seconds and long-term forgetting on the order of days – this is plausibly caused by depression of excitatory synapses or potentiation of inhibitory ones (Lim et al., 2015).

Previous modeling work on recognition memory used a predesigned architecture and plasticity rule and both empirical and analytic evaluation of performance (Androulidakis et al., 2008; Bogacz and Brown, 2003; Norman and O’Reilly, 2003; Sohal and Hasselmo, 2000). An emerging approach uses a machine learning technique known as “meta-learning,” or “learning how to learn” (Confavreux et al., 2020; Thrun and Pratt, 2012), which uses optimization tools to rapidly search for mechanisms that artificial neural networks can use to solve a learning/memory task. In contrast to hand-designed models, meta-learning enables unbiased exploration of a large family of architectures and plasticity rules. Importantly, it is possible to impose constraints that ensure biological plausibility (Bengio et al., 1991). For example, given a network architecture and a family of biologically plausible plasticity rules with tunable parameters, the meta-learning algorithm can search for the optimal parameters for memorizing a set of inputs.

In this work, we consider a family of models that recognize previously experienced stimuli and, importantly, learn and operate continuously without separate “learning” and “testing” phases. We investigate a feedforward network architecture with continual Hebbian plasticity in its synaptic weights. Parameters governing plasticity and other network parameters are meta-learned using gradient descent to optimize the continual familiarity detection process. To isolate synaptic plasticity as the unique memory mechanism, we avoid recurrent connectivity that could store memory through maintained neuronal activations. This architecture, unlike recurrent networks, generalizes naturally over a range of repeat intervals even if trained on a single interval. We show that an anti-Hebbian plasticity rule (co-activated neurons cause synaptic depression) enables repeat detection over longer intervals than a Hebbian rule, and this is the solution most frequently found by meta-learning. This rule leads to experimentally observed features such as repetition suppression in the hidden layer neurons. Critically, the capacity of these networks remains constant over time, so they can be continually fed new inputs with no reduction in steady-state memory performance.

## Results

### Continual familiarity detection task

We consider a continual familiarity detection task (Fig 1a) in which a stream of stimuli is presented to a network. With probability 1 − *p*, the stimulus at time *t* is chosen as a randomly generated *d*-dimensional binary vector ***x***(*t*), where each component is either +1 or −1 (note that for sufficiently large *d*, spurious chance repeats are extremely unlikely). With probability *p*, the stimulus is a copy of the stimulus presented *R* time steps ago, so that ***x***(*t*) = ***x***(*t* − *R*). However, we ensure that a stimulus is repeated at most once so, if ***x***(*t* − *R*) is already a repeat, i.e. ***x***(*t* − *R*) = ***x***(*t* − 2*R*), a new ***x***(*t*) is generated. As a result, the fraction of novel stimuli, which we call *f*, is not equal to 1 − *p*, but rather 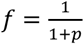. We use 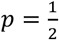, so 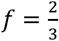 The output of the network should be *y*(*t*) = 0 if ***x***(*t*) is novel and *y*(*t*) = 1 if it is familiar, i.e. has appeared previously.

**Figure 1.**
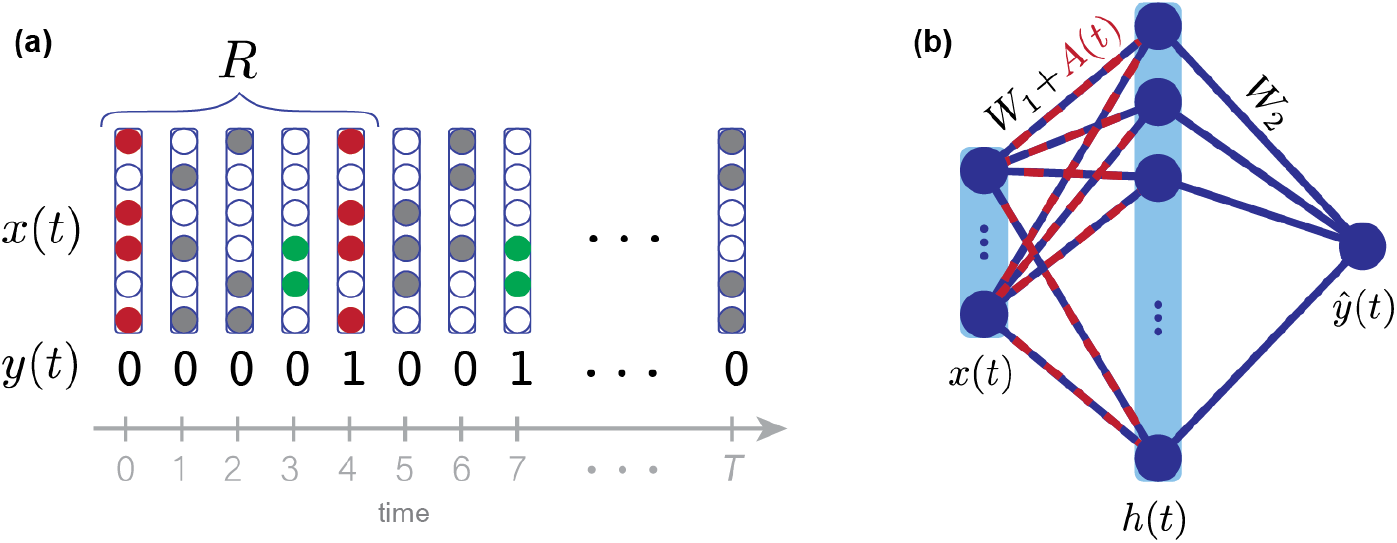
Continual familiarity detection task and HebbFF model. (a) The continual familiarity detection task. Given a continual stream of stimuli *x*(*t*), the desired output is *y*(*t*) = 1 if the stimulus has appeared previously and *y*(*t*) = 0 otherwise. For a given dataset, repeat stimuli always appear at an interval *R* after their first presentation. Although the task is continual, for the purposes of network training we use a finite-duration trial of length *T* ≫ *R*.(b) The HebbFF network architecture. A feedforward layer is endowed with ongoing Hebbian plasticity, the parameters of which are optimized using stochastic gradient descent. The hidden units are linearly read out to produce the network’s estimate of familiarity 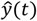.

The accuracy of the network (*P*_*correct*_, the probability of correctly responding to a stimulus) depends on two factors: the true positive rate (*P*_*TP*_, the probability of correctly reporting a repeated stimulus as “familiar“), and the false positive rate (*P*_*FP*_, the probability of incorrectly reporting a novel stimulus as “familiar“). These two factors are weighted by the fraction of novel stimuli *f*, so that *P*_*correct*_ = (1 − *f*) *P*_*TP*_+ *f*(1 − *P*_*FP*_). Through our choice of loss function (*Online methods*), we are effectively training the networks to maximize accuracy, so the “chance” level performance is *f* (for 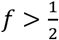), which a network can achieve by reporting all stimuli as novel (*P*_*TP*_ = *P*_*FP*_ = 0).

In our paradigm, a given dataset has a single repeat interval *R*, which differs slightly from previously studied experimental paradigms (Brady et al., 2008; Meyer and Rust, 2018). However, we evaluate performance on multiple datasets with various values of *R*. For testing, this is analogous to evaluating a single dataset with multiple repeat intervals and computing accuracy for each interval separately. We use this approach because it allows us to test generalization by training on one value of *R* and testing on others. It also allows us to train the network to its maximal capacity by gradually increasing *R* during “curriculum training”, and simplifies analytic calculations.

We begin by considering familiarity detection for uncorrelated stimuli, but, in later sections, we generalize to a task that requires simultaneous familiarity detection and binary classification, and to a dataset of real-world object images.

### HebbFF network architecture

To investigate the effectiveness of synaptic plasticity for solving this task, we use a feedforward neural network with a single hidden layer and, to implement the memory function, activity-dependent ongoing Hebbian plasticity (HebbFF) (Fig 1b). We purposefully do not include any recurrent connections to ensure that memory cannot be stored through persistent neuronal activity, thus isolating synaptic plasticity as the only possible memory mechanism.

In the HebbFF network, a group of hidden layer neurons with firing rates given by an *N*-dimensional vector ***h***(*t*), receives a *d*-dimensional input ***x***(*t*). The variable *t* indexes stimulus presentations which occur sequentially, so we refer to it as “time”. The input to each hidden-layer neuron is weighted by its corresponding synaptic strength and then transformed into a hidden-layer firing rate through a nonlinear activation function *σ*(*·*). The synaptic strength between the postsynaptic neuron with rate *h*_*i*_(*t*) and the presynaptic neuron carrying the input *x*_*j*_(*t*) is the (*i*, *j*) component of an *N*-by-*d* matrix that is the sum of a fixed matrix ***W***_1_ and a plastic matrix ***A***(*t*). Thus, the firing rate of the hidden layer is given by

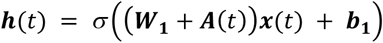

where *σ*(*·*) is the logistic function applied element-wise and ***b***_1_ is a vector representing baseline currents into the hidden layer. The plastic matrix ***A***(*t*) is updated at every time step: its (*i*, *j*) component decays by a factor 0 < *λ* < 1 and is incremented by a Hebbian product of the pre- and postsynaptic activities, *h*_*i*_(*t*)*x*_*j*_(*t*). A plasticity rate parameter – *∞ < η < ∞* controls the sign and magnitude of this increment. In matrix form, the synaptic update rule is then

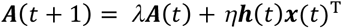

Finally, the output of the network 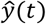 is a linear readout of the hidden layer and, since the target *y*(*t*) is binary, we bound the readout with the logistic function,

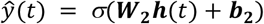

The response of the network is taken to be familiar if 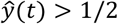 and novel otherwise.

To construct the network, we use backpropagation through time (BPTT) to “meta-learn” the parameters ***W***_1_, ***b***_**1**_, ***W***_**2**_, ***b***_**2**_, *λ*, *η*, which are fixed once training is completed (*Online methods*). The continual familiarity detection task – the “learning” task – is then performed exclusively by the ongoing synaptic dynamics of ***A***(*t*), determined by the fixed parameters. These dynamics are a biologically plausible mechanism for solving the continual memory task, but BPTT is simply used as an optimization tool to find suitable parameters of the network.

### HebbFF generalizes across datasets and repeat intervals

As a benchmark for comparing HebbFF performance, we first train a long short-term memory (LSTM) network (Hochreiter and Schmidhuber, 1997) – a recurrent neural network (RNN) architecture well-suited for memory performance – on the continual familiarity detection task. Unlike HebbFF, which stores its input history in the plastic synaptic matrix ***A***(*t*), an RNN uses ongoing neuronal activity.

If we train the RNN using a single dataset with *T* = 500 image presentations (*Online methods*) and a repeat interval of *R* = 3, it successfully learns the training set, but entirely fails to generalize to new test sets with the same *R* (Fig. 2a). This is because the RNN learns to perform classification, not familiarity detection. To fix this, we use an “infinite data” approach in which we generate a new dataset for every iteration of BPTT, each with the same value of *R* = 3. Trained in this way, the RNN now generalizes “in-distribution” across datasets with *R* = 3 (i.e. to datasets drawn from the same distribution as the training data, which is parameterized by *R*), but fails to generalize “out-of-distribution” to data with any other value of *R* (i.e. to datasets from a different distribution) (Fig 2b). The same result holds if we train another RNN with *R* = 6 (Fig 2b). We can further train the RNN with items spaced at intervals of both *R* = 3 and *R* = 6 (i.e. the value of *R* is chosen randomly for each familiar stimulus rather than being fixed). While the network can interpolate between the trained values, it does not extrapolate well to larger or smaller ones (Fig 2c). Although it is likely possible to train the RNN to perform well for multiple values of *R* by using more complex training schedules, we believe that poor out-of-distribution generalization is a bottleneck of the RNN approach.

**Figure 2.**
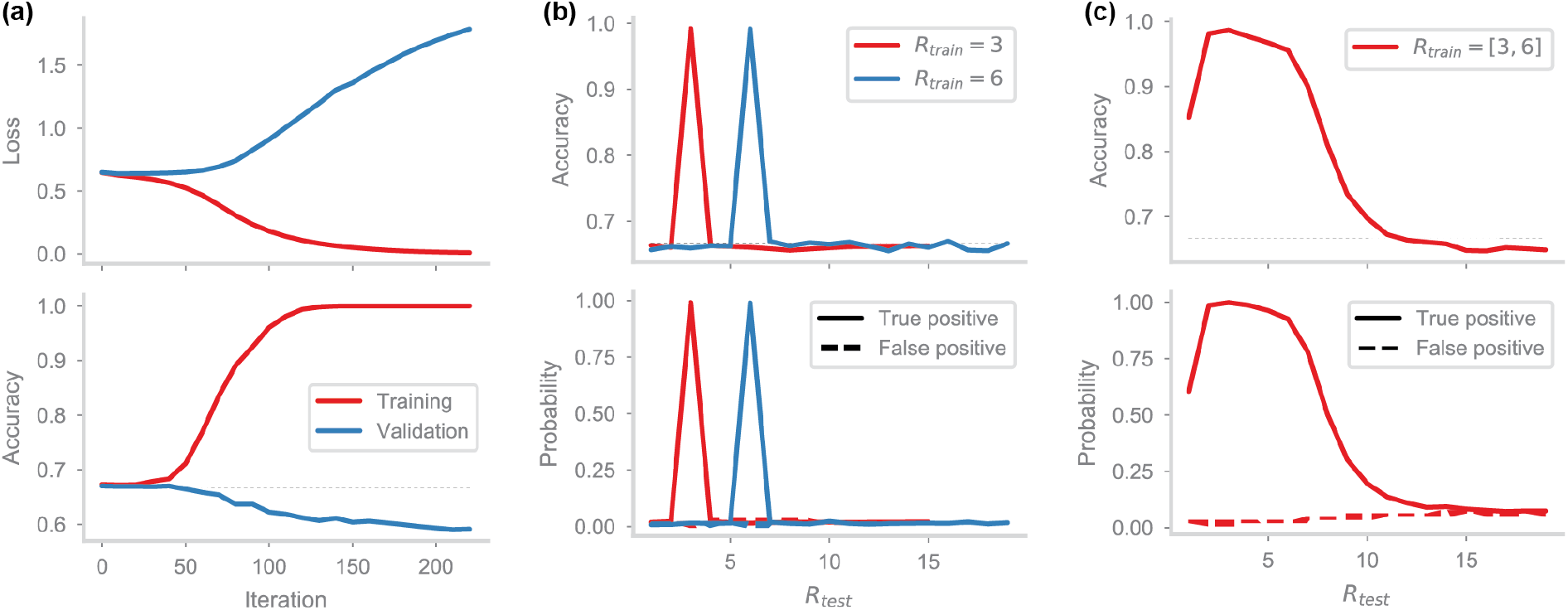
RNN performance on continual familiarity detection. (a) Training an LSTM (*d* = 100 input dimension, *N* = 100 recurrent units) on a single dataset of a familiarity detection task (*T* = 500 stimulus presentations, repeat interval *R* = 3). Although the loss (top) approaches zero and accuracy (bottom) approaches 1 on the training dataset (red curves), performance on a validation dataset (blue) with the same parameters fails to generalize even when tested in-distribution with the same *R*. (b) Training the RNN using “infinite data.” New datasets, each with *R* = 3, are generated at every epoch of training (red). Accuracy (top), as well as true positive and false positive probabilities (bottom) is shown as a function of the repeat interval on validation datasets. The same is repeated with another RNN using *R* = 6 (blue). The RNNs perform well in-distribution on datasets with the same repeat interval as used during training, but fail to generalize out-of-distribution to other repeat intervals. (c) Training the RNN with “infinite data,” using datasets with both repeat intervals *R* = 3 and *R* = 6. The RNN interpolates between the intervals – performance is high when tested on repeat intervals 3 ≤ *R* ≤ 6 – but fails to extrapolate. Performance quickly drops for longer repeat intervals, and even for shorter ones.

In contrast, the HebbFF network exhibits both in-distribution and out-of-distribution generalization. Even when trained on a single dataset with a fixed repeat interval *R*, the network generalizes not only to new test sets with the same *R*, (Fig 3a) but even to those with different *R* values. Trained with “infinite data” (the scheme we use in general), HebbFF generalizes to datasets with smaller and even larger *R* values (Fig 3b). If we match the number of dynamic variables rather than the number of hidden neurons, HebbFF still shows superior generalization compared to the RNN (Fig S1). This qualitative difference in performance suggests that Hebbian plasticity provides a powerful inductive bias in the form of a more “natural” mechanism of memory for the purpose of familiarity detection.

**Figure 3.**
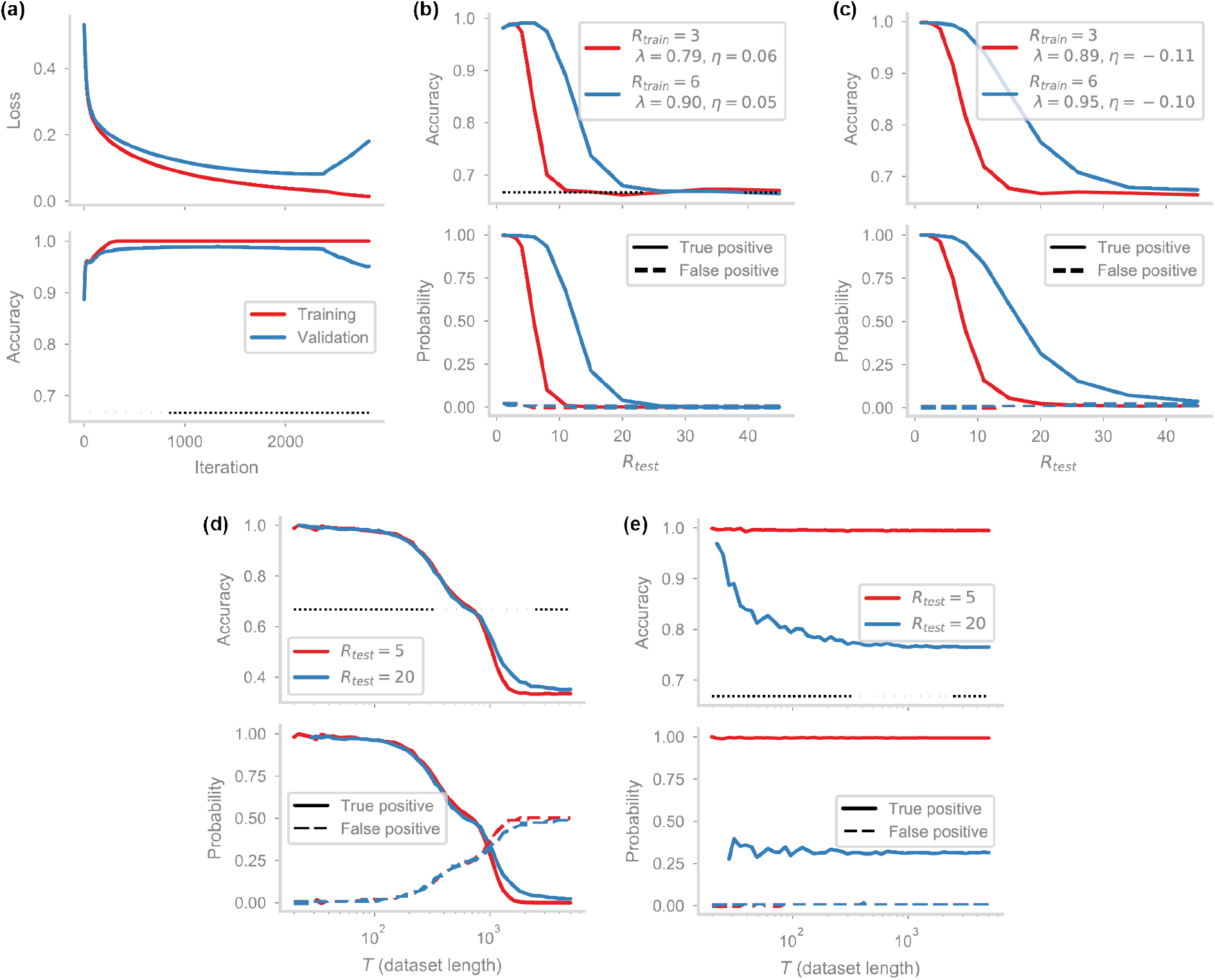
Hebbian vs. anti-Hebbian plasticity and continual operation. (a) Training the HebbFF network (*d* = *N* = 100), as in Fig 2a. Both training and validation loss decrease, and accuracy increases, for a single instance of the dataset with *R* = 3, indicating in-distribution generalization. Over many iterations, however, overtraining occurs due to the use of a single dataset, increasing the final validation loss. (b) HebbFF network trained with “infinite data” as in Fig 2b (*R* = 3, red; *R* = 6, blue) shows not only in-distribution generalization to any dataset with *R* = 3 (*R* = 6, resp.), but also out-of-distribution to datasets with any smaller *R* and some larger *R*’s. (C)HebbFF with a different initialization converges to a qualitatively different solution with a negative learning rate η, an anti-Hebbian learning rule, in contrast to the Hebbian solution in (b). The anti-Hebbian solution shows generalization performance over a larger range of *R* values than Hebbian. (d) Model from (Bogacz and Brown, 2003), evaluated on the continual familiarity detection task, varying the length *T* of the trial. Accuracy (top) is near-perfect regardless of the repeat interval *R* (blue vs. red curve) until the model reaches its capacity (*P*\* ≈ 100 for network size *d* = *N* = 100) because the model reliably stores the first *P*\* patterns. Accuracy rapidly drops below chance for *T* > *P*\* as the model begins to report familiar stimuli as novel (see Fig S2b). (e) HebbFF network operates continuously, as its accuracy (top) is consistent with the generalization curve from (c), with near-perfect performance for *R*_test_ = 5 and above 80% for *R*_test_ = 20 for any trial length. True and false probabilities (bottom) are better representations as accuracy (top) is artificially higher for small *T* due to the low proportion of familiar stimuli.

The generalization performance of HebbFF is due to the fact that the memory representation of an item does not change over time, other than being scaled by a factor. A stimulus ***x***(*t*) is initially stored as the outer product of ***h***(*t*) and ***x***(*t*), multiplied by the plasticity rate *η*. The plastic component of the connectivity matrix also contains terms arising from previously stored memories which, for the purposes of this particular stimulus, act as additive noise *ε*:

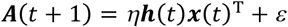

As subsequent stimuli are presented, the representation of ***x***(*t*) maintains the same form, so that *k* time steps later it is still stored as the outer product of ***h***(*t*) and ***x***(*t*), scaled by a factor λ^*k*^:

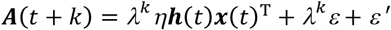

where further additive noise *ε*’ arises from stimuli presented after ***x***(*t*).

Unlike HebbFF, RNNs are poor at generalizing across intervals *R* because the dynamics of their units allow the memory representation of a stimulus to change arbitrarily over time. The RNN only generates the appropriate representation at the time when a query is expected, namely after a delay equal to the value of *R* used during training. This makes it difficult to generalize across intervals.

### Anti-Hebbian outperforms Hebbian plasticity

The plasticity rate *η* in HebbFF can be positive or negative, resulting in either Hebbian or anti-Hebbian plasticity. For the Hebbian solution with *η* > 0, synapses are potentiated in response to a stimulus. When it is repeated, the hidden layer activity is higher than for a novel stimulus due to the increased strength of the synapses storing the memory. For anti-Hebbian plasticity, *η* < 0, synapses are depressed when a memory is stored. In this case, the hidden layer activity is lower for a familiar stimulus than a novel stimulus, which is consistent with experimental results of repetition suppression (Grill-Spector et al., 2006; Meyer and Rust, 2018; Xiang and Brown, 1998). Furthermore, the meta-learning algorithm is more likely to converge to the anti-Hebbian solution, especially when trained with a relatively large repeat interval, even if the initial value of *η* is positive, and almost always does so when the initial value is negative.

Interestingly, anti-Hebbian plasticity enables successful familiarity detection over considerably longer intervals than a Hebbian rule (Fig 3c). To understand this, note that the memory of a stimulus is degraded in two ways: additional plasticity events obscure existing memories, and the plastic weights decay over time. With an anti-Hebbian plasticity rule, the hidden layer activation ***h***(*t*) is close to zero for a familiar stimulus due to repetition suppression. As a result, the plasticity update *η****h***(*t*)***x*** (*t*)^*T*^ when the stimulus is repeated is negligible – as if a stimulus was not presented at that time step. This effectively reduces the number of plasticity events, reducing the disruption of existing memories. As a secondary effect, the smaller number of plasticity events allows a larger *λ*(smaller decay rate) to be used while still controlling the amplitude of plastic weights. This slower decay rate further extends the lifetime of the memory. Due to their superior performance and consistency with experimental results, we only consider anti-Hebbian solutions throughout the following sections.

### HebbFF learns continually without catastrophic forgetting

Previous modeling work using anti-Hebbian plasticity mechanisms for familiarity detection (Bogacz and Brown, 2003) focused on a paradigm used in classic studies of recognition memory (Standing, 1973) in which subjects are serially presented an entire dataset and later asked to identify which stimulus is familiar in a two-alternative-forced-choice (2AFC) test. Analogously, this previous modeling work used explicit “learning” and “testing” phases and demonstrated an impressive capacity for recognition memory (Bogacz and Brown, 2003) (Fig S2a). When evaluated on the continual memory task that we use, the Bogacz-Brown model has near-perfect performance if the number of stimuli *T* in the dataset is smaller than the model’s capacity *P*^*^, independent of the value of the repeat interval *R* (Fig 3d). That is, the model successfully stores all *T* < *P*^*^ stimuli. As the dataset size *T* increases, however, the model performance declines due to catastrophic interference (Fig 3d, S2b; *Online methods*). To store additional memories, the old memories must be removed by resetting the synaptic weights.

In contrast, the HebbFF model operates continually rather than using separate learning and evaluation phases. Its performance is independent of the length of the dataset, and it can operate continuously without any need to reset the synaptic weights. For example, a HebbFF network trained with *R* = 5 operates at near-perfect performance irrespective of the duration of the trial *T* when tested with *R* = 5 (Fig 3e). Similarly, when tested with *R* = 20, it operates continually at near 80% accuracy (Fig 3e), as expected from the generalization curve in Fig 3c (note that for small *T* the accuracy (Fig 3e, top, blue) is transiently elevated because the fraction of novel stimuli is more than 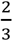). In other words, the model has a 3 moving window in time within which it can successfully detect a familiar stimulus, and it forgets old stimuli gracefully without suffering from catastrophic interference.

### A uniform readout

Up to now, the readout of the hidden layer, 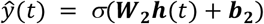, has involved a trained matrix W_**2**_. Because the inputs to the network are uniformly random, there is no reason to believe that, as far as the output is concerned, one hidden unit would be statistically different than any other (although they may still be statistically dependent). This observation leads us to restrict ***W***_**2**_ to be a scaled 1-by-*N* matrix of ones, ***W***_**2**_ = *α*_2_[1,…,1], where *α*_2_ is a trained scalar. Similarly, we restrict ***b***_1_ = *β*_1_[1,…,1]^*T*^. We verified that performance is not affected by this choice of output weights (Fig S3a), the distribution of 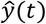 for familiar and novel stimuli is the same (Fig S3b), the readout vector ***W***_**2**_ and the bias term ***b***_1_ have similar features (Fig S3c), and anti-Hebbian plasticity is still the preferred form of plasticity. We use this uniform readout throughout the remainder of our studies unless stated otherwise.

### Storage and readout mechanisms

In the HebbFF network, the hidden layer plays a dual role. On the one hand, it must produce a reliable familiarity signal for the readout to decode. On the other, it must create a robust representation of the input stimulus during the Hebbian plasticity update. The hidden activity is controlled by the fixed parameters ***W***_1_ and ***b***_1_, as well as the plastic matrix ***A***(*t*). Here, we investigate how ***W***_1_, ***b***_1_ and ***A***(*t*), impact these two aspects of the familiarity detection task.

Networks trained with larger *R* have sparser hidden unit activity (Fig 4a-c): the sparser the activity, the less plasticity is evoked, and thus the longer memories can be retained without overwriting. In the limiting case we might expect that exactly one neuron is active for a novel stimulus and none are active for familiar stimuli. Associated with this increased sparsity in activity, ***W***_1_ is also sparser for larger *R* (Fig 4d-f).

**Figure 4.**
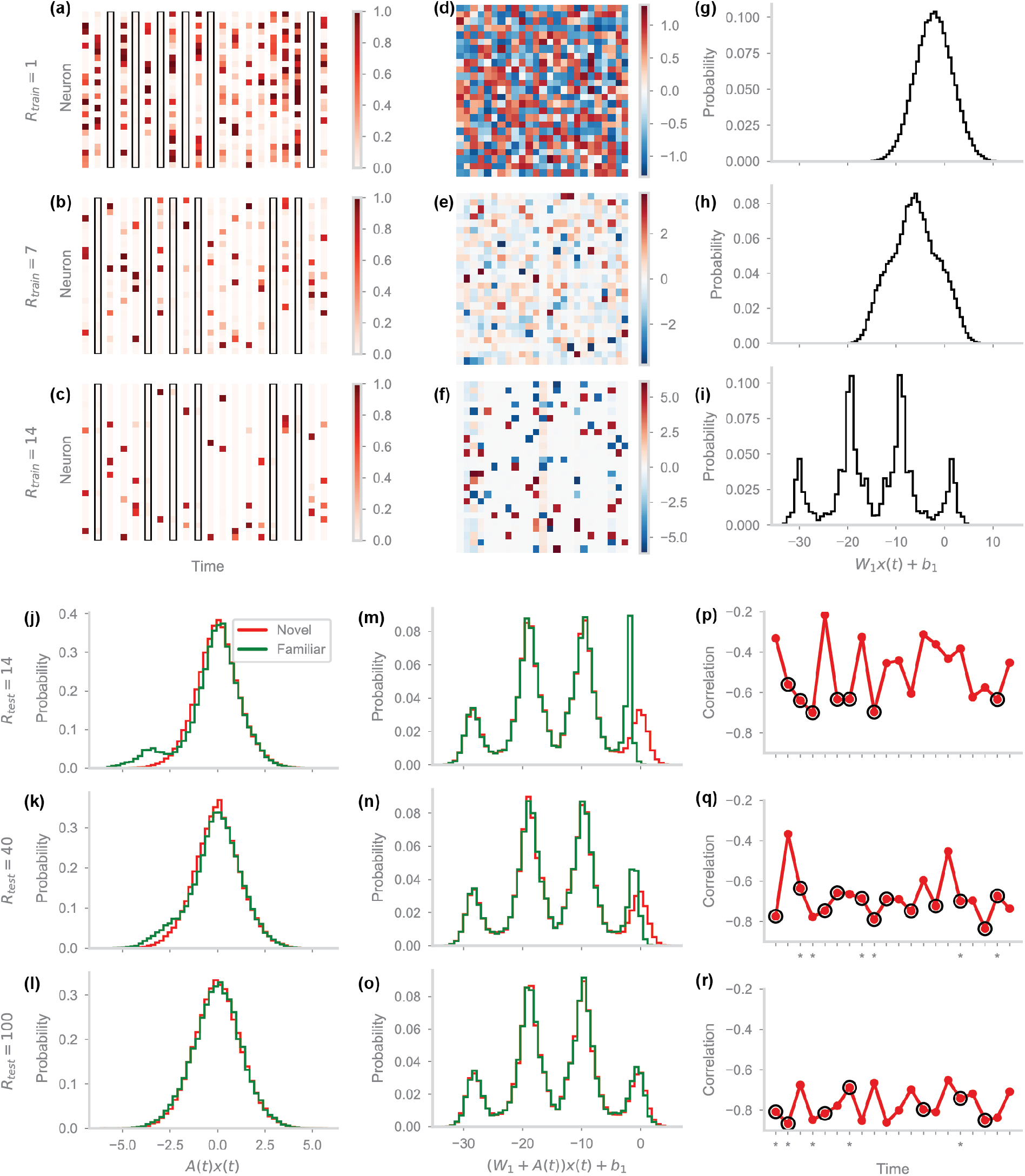
Storage and readout mechanism. (a-c) Hidden layer activity ***h***(*t*) over 20 consecutive timesteps for networks with input dimension *d* = 25 and *N* = 25 hidden units, trained on datasets with *R* = 1, 7, or 14, respectively. Familiar stimuli (black rectangles) cause silencing i.e. repetition suppression of hidden layer activity. Activity for novel stimuli becomes sparser for networks trained with larger *R*. (d-f) Static weight matrix ***W***_1_ of the networks from (a-c). The weight matrix becomes sparser, and individual weight magnitudes increase for networks trained with larger *R*, enabling more sparse activity in the hidden layer for novel stimuli. (g-i) Distributions of input current into the hidden layer due to the static component of the synapses, i.e. the matrix ***W***_1_ and bias ***b***_1_, for the networks from (a-c). We do not distinguish familiar and novel stimuli since the current due to the static component is the same regardless of novelty. For networks trained with larger *R*, the distribution becomes multi-modal, with the number of modes equal (approximately) to the number of high-magnitude values per row of ***W***_1_, plus one. Due to the bias, only the rightmost mode has the potential to produce firing rates that are significantly above zero. (j-l) Distributions of input current into the hidden layer due to the plastic component of the synapses, i.e. the matrix ***A***(*t*), for novel (red) and familiar (green) stimuli. We only consider the trained network from (c,f,i) and evaluate its behavior on test sets with *R* = 14, 40, or 100, corresponding to perfect, intermediate, and chance accuracy. The large central mode occurs due to stored stimuli uncorrelated with the input stimulus ***x***(*t*). In the novel case, the input is uncorrelated with all the stored stimuli by definition, and thus there is only one mode. Similarly, in the familiar case with a long delay interval *R* = 100 the stored stimulus has decayed sufficiently that its signal is lost. In the case of familiar stimuli presented at shorter delay intervals, *R* = 14 or 40, there is an additional mode due to the correlation between the input ***x***(*t*) and its copy ***x***(*t* − *R*) previously stored in the plastic matrix ***A***(*t*). (m-o) Distributions of the total input current into the hidden layer on test sets with *R* = 14, 40, or 100. Only the values above zero cause high firing rates after applying the logistic sigmoid nonlinearity. Since all the input currents are low for familiar stimuli (green) for small values of *R*, there is repetition suppression. (p-r) Correlation between the input current into the hidden layer from static and plastic synapse components at each of 20 consecutive timepoints. Asterisks indicate output response errors. For sufficiently small *R*, the input currents are more anti-correlated for familiar stimuli (black circles) than for novel. Combined with the distributions of input currents, this indicates that the units receiving positive input current from the static synapses receive negative input current from the plastic synapses.

To isolate the effect of ***W***_1_ on hidden unit activity, we compute a histogram of the input current into the hidden layer due to the non-plastic synapses, *W*_1_***x***(*t*) + ***b***_1_, across units and across time (Fig 4g-i). As *R* increases, the distribution becomes multi-modal. In general, the number of peaks in this distribution depends on the number of large-magnitude values of ***W***_1_ per row. Critically, due to the logistic function nonlinearity, only the rightmost peak in Fig 4i is large enough to elicit appreciable activity in the hidden layer. This peak drives the small number of hidden units that are significantly activated by a novel stimulus. In other words the ***W***_1_ matrix acts like a hash function to select a small subset of hidden units to store the memory of a given stimulus.

We next consider the effect of ***A***(*t*), focusing on the network trained to maximum capacity (Fig 4c,f,i) (see next section). For a novel stimulus, the distribution of the input current due to the plastic synapses ***A***(*t*)***x***(*t*) is unimodal and symmetric about zero (Fig 4j-l). For a familiar stimulus, however, there is an additional peak at approximately *λ*^R−1^*ηd*. This peak is due to the dot product of the input vector ***x***(*t* −*R*) (stored in the matrix ***A***(*t*) as *λ*^R−1^*η****h***(*t*−*R*)***x***(*t*−*R*)^T^), and the familiar input vector ***x***(*t*) = ***x***(*t*−*R*). Importantly, the neurons that exhibit this behavior are the same neurons that were active due to ***W***_1_ when the stimulus was novel. This implies that, in addition to selecting a subset of neurons for storage of a memory, ***W***_1_ also allows the system to probe those same neurons during recall. Thus, ***W***_1_ serves as an addressing function during memory recall.

Finally, the total hidden layer input current is the sum of these two components, (***W***_1_ + ***A***(*t*))***x***(*t*) + ***b***_1_ (Fig 4m-o). Comparing Fig4i and Fig4o, we see that the large central symmetric mode of the ***A***(*t*)***x***(*t*) distribution does not significantly impact the total hidden layer input current. Rather, the familiarity signal arises because the smaller peak of the ***A***(*t*)***x***(*t*) distribution pushes the rightmost peak of the ***W***_1_***x***(*t*) + ***b***_1_ distribution below zero (Fig 4m). Anti-correlation between the two input currents for familiar stimuli (Fig 4p-r) indicates that this shift is caused by the input current from the plastic component of the synapse cancelling the input current from the fixed component, resulting in lower activation, i.e. repetition suppression.

### Curriculum training and memory capacity

A randomly-initialized HebbFF network is typically unable to find a solution if trained with a large value of *R* (e.g. a network with *d* = 50, *N* = 100 rarely converges if trained on *R* > 10). Instead, we use a curriculum training procedure to bootstrap the optimized solution. First, the network is trained on data with *R* = 1, using the “infinite data” regime. Once the accuracy is above 99%, *R* is incremented by one and training continues on data with *R* = 2. This process continues until *R* becomes large enough that the network cannot find a solution with accuracy above 99%, i.e. if *R* is not incremented for at least 2 million iterations (Fig 5a). We thus define the memory capacity *R*_max_ as the largest value of *R* for which the familiarity detection accuracy is above 99%.

**Figure 5.**
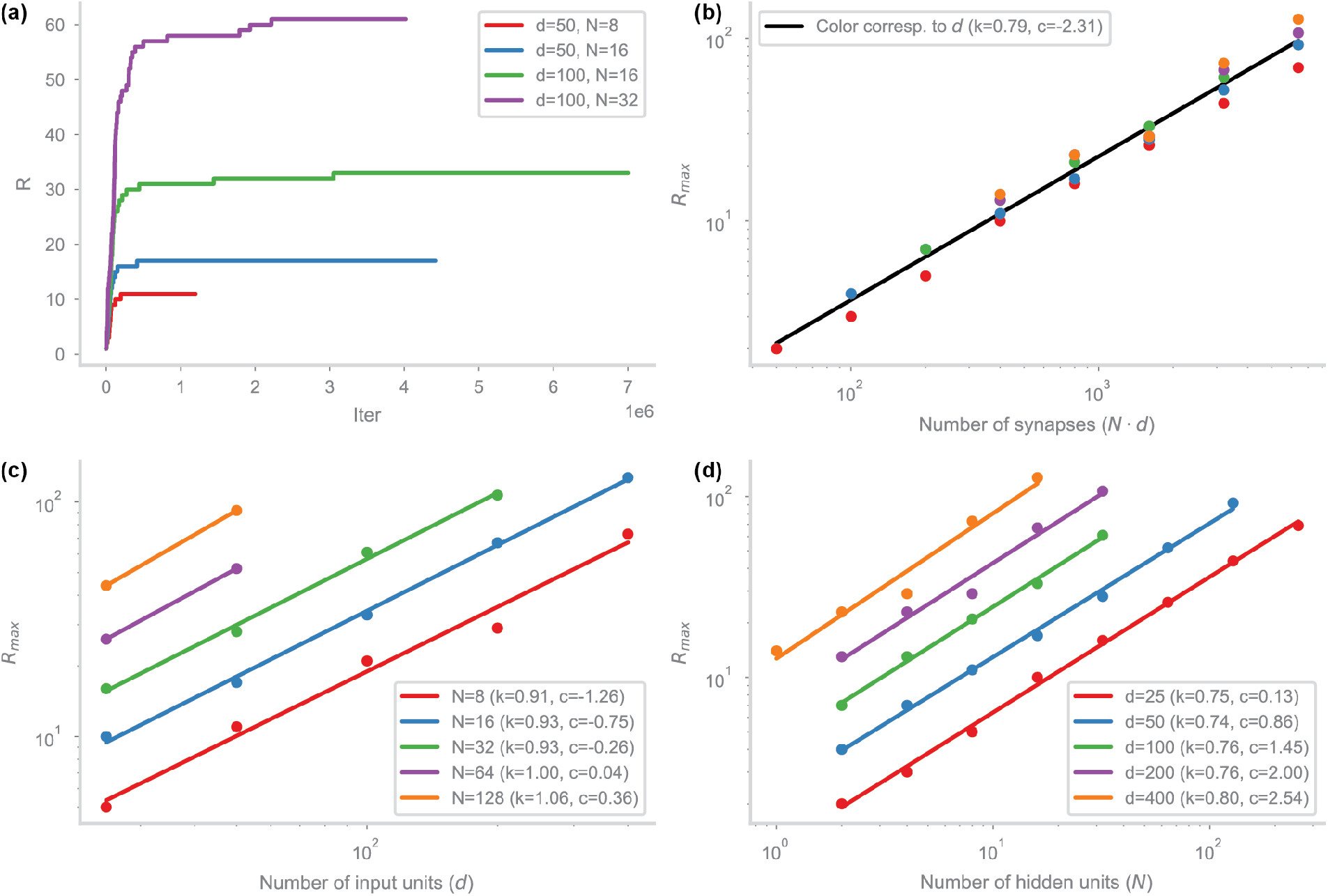
Curriculum training and empirical capacity. (a) The value of *R* used over the course of curriculum training for four different network sizes. *R* is incremented once the network achieves an accuracy > 0.99. Training is considered converged when the value of *R* is not incremented for at least 1 million iterations. (b) The final value of *R* after curriculum training (i.e. network capacity) as a function of the number of plastic synapses in the network, plotted on a log-log scale. The color of the points corresponds to the number of input units, colors from panel (d). The least-squares fit (black line, slope *k*, bias *c*) indicates that the empirical network capacity scales sub-linearly with the number of synapses. (c) Capacity as a function of the input dimension *d* for various hidden layer sizes *N*. (d) Capacity as a function of the hidden layer size for various input dimensions *d*. Capacity primarily depends only on the number of synapses, rather than on the hidden or input layer sizes.

We curriculum-train networks of different sizes and plot the capacity *R*^max^ for each one (Fig 5b). For consistency and ease of training, we restrict the networks to the anti-Hebbian solution and use the uniform readout. We find that the capacity depends primarily on the number of synapses, rather than on the number of pre- or postsynaptic neurons (Fig 5c,d), consistent with previous familiarity detection results (Bogacz and Brown, 2003). To estimate the scaling, we compute a linear least-squares fit of log(*R*^max^) as a function of log(*Nd*). Empirically, we find that the capacity of the network scales as

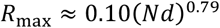

which is sublinear in the number of plastic input synapses to the hidden layer, *Nd*.

The model of Bogacz and Brown (Bogacz and Brown, 2003) for the non-continual task has a capacity that is linear in the number of synapses. To determine whether the difference between the empirical performance of HebbFF and the Bogacz-Brown model reflects a fundamental limitation in the feedforward architecture, we developed an idealized version of the model (Fig 6a) that we could study analytically (*Online methods*).

**Figure 6.**
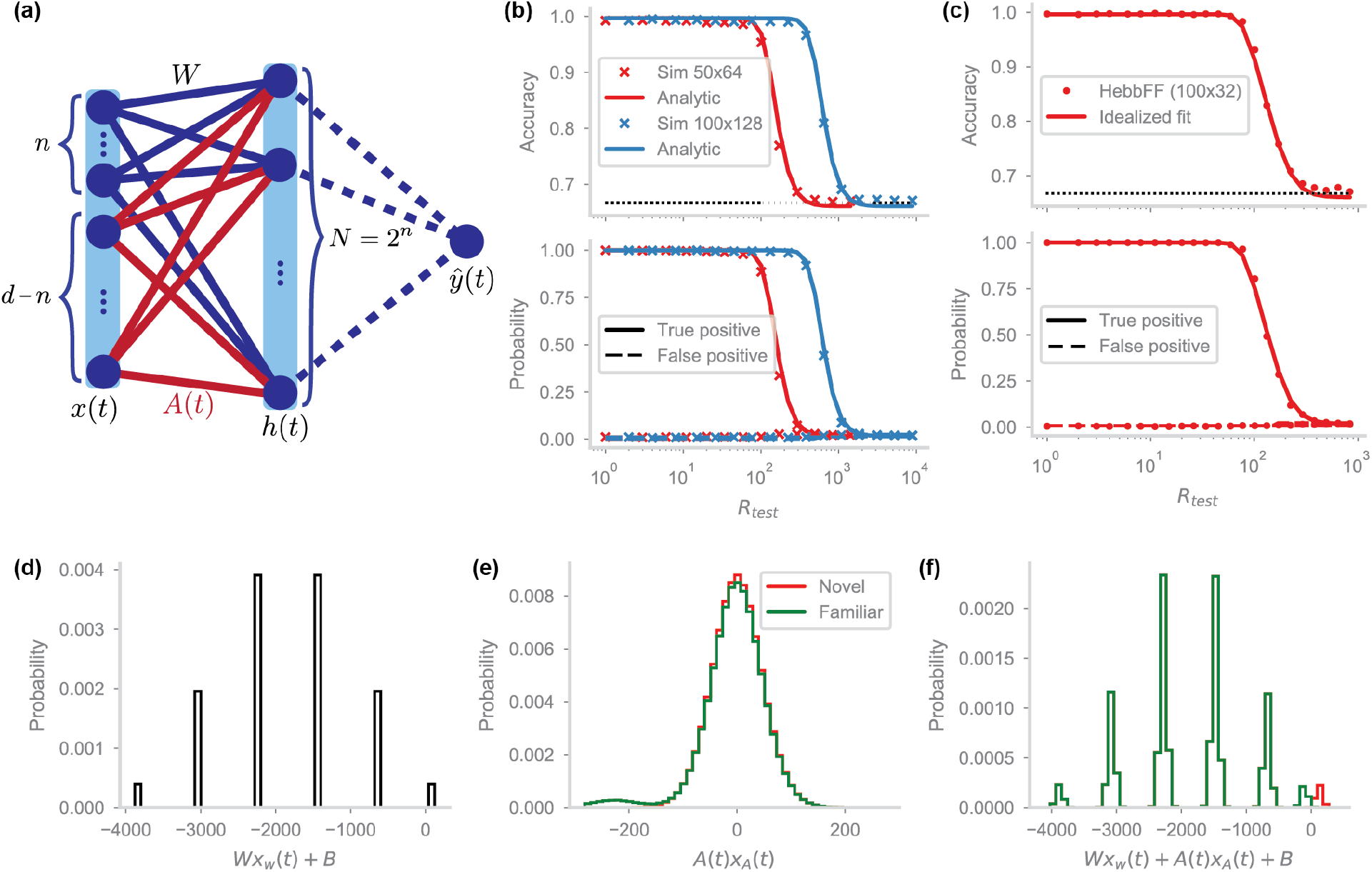
Idealized model. (a) The idealized HebbFF network architecture. In contrast to the original HebbFF network with a single effective matrix ***W***_1_ + ***A***(*t*), the input ***x***(*t*) is effectively split into two sections of size *n* and *D* = *d* − *n* that serve as inputs into separate static and plastic synaptic matrices ***W***_1_ and ***A***(*t*), respectively (*Online methods)*. The hidden layer size is *N* = 2^*n*^. The readout unit outputs 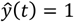 whenever any of the hidden units is active. (b) The analytic calculation of network performance (solid line) matches simulation results for the idealized network (x’s), shown for two different network sizes (red, blue). (c) A least-squares fit of the analytic performance curve of the idealized network to a trained HebbFF network of the same size for two network sizes. The idealized network has similar performance to the HebbFF model if its decay rate and bias are scaled appropriately: λ ≈ 0.986, *b*_1_ ≈ −4.771 (for all units) for *d* = 200, *N* = 32, and λ ≈ 0.993, *b*^1^ ≈ −4.771 for *d* = 200, *N* = 32. (d-f) Same as Fig 4(I,j,m), but for the idealized network (*D* = 400, *N* = 32, *R* = 300).

We noted above that the limiting behavior of the network at maximum capacity appears to have ***W***_1_ activate just a single unit for memory storage. We build this limiting behavior into the idealized model through a specific choice of ***W***_1_ and ***b***_1_, set by design rather than through a training procedure. Specifically, we use the first *n* ≪ *d* components of ***x***(*t*) as an identifier by choosing the first *n* columns of ***W***_1_ so that a unique hidden unit is activated by each possible *n*-bit combination of these components, and set the remaining columns of ***W***_1_ to zero. To simplify the model, we do not allow plasticity to operate on the inputs from these bits and set the first *n* columns of ***A*** *t* to zero (Fig 6a). This isolates the “hashing” function of the fixed matrix from the memory storage. Furthermore, instead of a sigmoid nonlinearity for the hidden units, we use a Heaviside step function Θ(·). Thus, the hidden layer in the idealized model is governed by

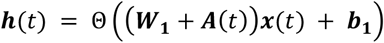

For the nonzero entries of ***A***(*t*), plasticity is the same as in the trained model. However, because the Heaviside function does not depend on the scale of the input, we can set the plasticity rate to η = −1 without loss of generality. The optimal synaptic decay rate λ can be computed analytically. Finally, a stimulus is considered familiar if all hidden unit activities are identically zero, and novel otherwise (*Online methods*).

This idealized model exhibits qualitatively similar behavior to HebbFF. We can fit the analytic functional form of the true and false positive probabilities computed from the idealized model (Fig 6b) to the corresponding probabilities of HebbFF (Fig 6c). Furthermore, the histograms of inputs to the hidden layer are qualitatively similar: ***W***_1_***x*** *t* + ***b***_1_ has the same multimodal distribution with more prominent peaks in the middle (Fig 4i, 6d), a bimodal distribution of ***A***(*t*)***x***(*t*) with a large symmetric central peak and a smaller one corresponding to the familiarity signal (Fig 4j, 6e), and a similar distribution of the total input current (***W***_1_ + ***A*** *t*)***x*** *t* + ***b***_1_ (Fig 4o, 6f). From this, we conclude that the memory storage and readout mechanisms are analogous in the meta-learned HebbFF network and the idealized model.

Along with the true and false positive probabilities, the memory capacity of the idealized model can be computed analytically (*Online methods*). As in the Bogacz-Brown model (Bogacz and Brown, 2003), the capacity, as characterized by 99% accuracy, is proportional to the number of synapses *Nd*. There are several possible reasons for the discrepancy between this analytic capacity, as well as that of the Bogacz-Brown model, relative to the empirical capacity for HebbFF.

First, the idealized HebbFF model uses a dedicated set of synapses through the fixed ***W***_1_ matrix, and the Bogacz and Brown model selects the units that have the highest input current implicitly through inhibitory competition. Both of these are dedicated addressing functions for the hidden layer, but meta-learned HebbFF must multiplex this functionality with memory storage, leading to correlations between the hidden layer input currents from the plastic and fixed synapse components (Fig S4a).

In addition, replacing the logistic function with a Heaviside function means that familiar stimuli truly generate no plasticity in the idealized model, reducing overwriting at the cost of not reinforcing partially-decayed memories (Fig S4b-c). For the same reason, in contrast to HebbFF, the idealized model achieves maximal plasticity for any suprathreshold level of input to a hidden layer unit.

Finally, training the HebbFF model may lead to specialized solutions for small *d* and *N* that have better performance than that predicted by the asymptotic analysis. Similarly, training may not converge to the optimal solution for large *d* and *N* because it requires the use of very long repeat intervals *R*. This means the dataset size *T* must be very large to include a sufficient number of familiar examples, which may lead to practical issues such as vanishing gradients. Thus, the empirical capacity may scale sublinearly with the number of synapses because of over-performance at low *R*, under-performance at high *R*, or both.

### HebbFF recapitulates neural data from inferotemporal cortex

We next compare the optimized HebbFF model with experimental results. Meyer and Rust (Meyer and Rust, 2018) recorded neurons from the inferotemporal (IT) cortex of monkeys performing familiarity detection and compared the quality of two decoders in predicting behavior from neural data as a function of neural population size. The authors considered a “spike count classifier” (SCC) decoder, which amounts to comparing a simple average of neuronal firing rates to a threshold, as well as a Fisher linear discriminant (FLD), which instead considers a weighted average, with weights computed from the data (Meyer and Rust, 2018).

We perform a similar analysis. We first construct an FLD decoder of the hidden unit firing rates and rank the units in reverse order of their FLD readout weights (i.e. units with the most negative weights are top-ranked; *Online methods*). We then consider decoders that use increasingly larger subsets of hidden units, adding them according to their ranking (Meyer and Rust, 2018). As in the experimental data, performance saturates for the FLD and declines for the SCC readout beyond a certain number of decoded units (Fig 7a). This can be explained by the fact that some units do not provide a reliable signals of familiarity – in the network this is due to suboptimal training, and in the IT cortex possibly due to those neurons performing an unrelated task (see next section). Including them hurts performance of the SCC decoder, but since the FLD readout weight for these units is close to zero, they do not alter its familiarity detection performance.

**Figure 7.**
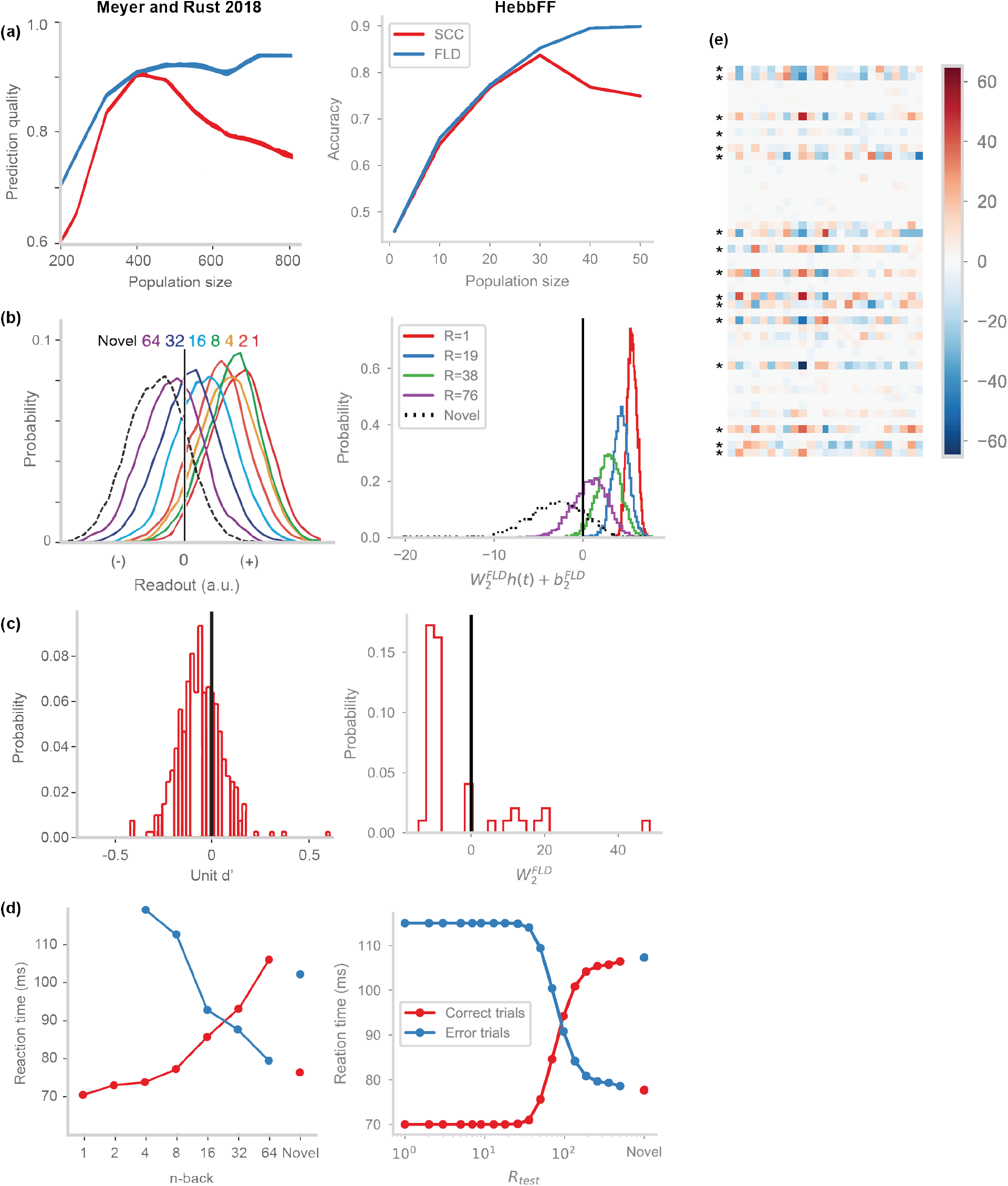
Comparison to IT cortex data. (a) Left: neurons from the IT cortex used to predict the behavioral outputs of a monkey performing continual familiarity detection, decoded using the Fisher linear discriminant (FLD, blue) or spike count classifier (SCC, red). Right: units from the hidden layer of a trained HebbFF network (trained with an unconstrained readout ***W***_**2**_ rather than uniform) used to detect familiarity obtained from SCC and FLD decoders. In both cases, the number of neurons/units available to the decoder was varied, added in order of increasing weight according to the FLD decoder. While the FLD decoder accuracy saturates, the SCC decoder accuracy peaks and begins to decline as more neurons/units are included in the decoder. (b) Distribution the FLD decoder output for IT cortex neurons (left) and HebbFF hidden units (right) for novel stimuli, and familiar stimuli at varying delay intervals. In both cases, the distribution shifts towards lower values as delay interval increases. For HebbFF, the distribution gets narrower for shorter delay intervals due to saturation in the hidden layer units. (c) Distribution of the FLD decoder weights for decoding IT cortex data (left) or HebbFF hidden unit activity (right). In both cases, the majority of output weights are negative, with some positive values. (C)Left: measured reaction time as a function of delay interval for correct and error trials (red, blue curves) in monkeys performing the continual familiarity detection task. Black lines indicate reaction times predicted using strength theory analysis. Right: HebbFF predicted reaction times using analogous strength theory analysis, using constants of proportionality from (Meyer and Rust, 2018) (*Online methods*). Both result in a qualitatively similar x-shaped pattern. Plots on the left side of (a-d) adapted from (Meyer and Rust, 2018). (e) The weight matrix ***W***_1_ of the HebbFF network trained on the augmented task, requiring simultaneous classification and familiarity detection. Hidden units split into “classification” and “familiarity” units, with classification units (marked with asterisks) having very strong input weights to overcome the noise from the plastic ***A***(*t*) matrix.

Comparing the experimental and model distributions of readout activity shows a qualitatively similar pattern for outputs to novel and familiar stimuli (Fig 7b). Both distributions shift toward smaller values as *R* increases, as the outputs for familiar stimuli approach those for novel. The fact that the distribution of outputs becomes narrower for HebbFF as *R* decreases, unlike in the data, may be due to repetition suppression causing hidden units to have near-zero responses for highly familiar (low *R*) stimuli, thus causing the readout distribution to cluster around its minimal value. On the other hand, biological neurons that exhibit repetition suppression may never be fully silenced – for example if it takes multiple repetitions to achieve maximal familiarity or if neurons are multiplexed with another task that requires a baseline level of activity. Furthermore, as in the data, the distribution of readout weights is biased towards negative values (Fig 7c).

Finally, using a similar “strength theory” analysis as in the experimental results (Meyer and Rust, 2018; Murdock, 1985), which suggests that reaction times are inversely proportional to the distance of the readout from the threshold, we can qualitatively reproduce the x-shaped reaction time curves seen in the data (Meyer and Rust, 2018). We used the same proportionality constant determined experimentally to compute network “reaction times” (Fig 7d). Overall, we find that the HebbFF model captures a number of features seen in the experimental results.

### Two subpopulations emerge in a classification-augmented task

IT cortex encodes object identity as well as familiarity (Lehky and Tanaka, 2016; Lueschow et al., 1994). To match this dual functionality, we augment familiarity detection with object classification. We first create a large pool of random vectors and randomly assign a binary label to each one. We then generate a familiarity detection dataset as before, except that each novel input is drawn from this pool (without replacement) rather than being generated anew. In addition to the scalar readout of familiarity, the network must now report the class of the stimulus through a second binary output. Critically, both outputs are read out from the same hidden layer activity (*Online methods*).

The augmented task could be solved by having all the neurons multiplexed to encode both familiarity and object identity. Alternatively, the neurons could split into two subpopulations, one of which detects familiarity and the other classifies objects (Rutishauser et al., 2015). We find that the HebbFF model converges to this second solution, an even split between familiarity and classifier units, as evident from inspecting the ***W***_1_ matrix (Fig 7e). Consistent with this, the capacity of the classifier-augmented HebbFF with 50 hidden units (*R*^max^ ≈ 13) is approximately the same as the original network with 25 units (*R*^max^ ≈ 14). In accord with this split, SCC decoder performance peaks in the split-task network when half of the top-ranked units are included (Fig S5d) because including units responsible for object identity but not familiarity degrades the familiarity readout. The other similarities to experimental results discussed in the previous section also hold for the task-augmented network (Fig S5).

### Familiarity detection of real images

To validate the HebbFF model in a more realistic scenario, we evaluate its performance on real-world object images. We consider the dataset used by Brady et al. to study familiarity detection in humans (Brady et al., 2008). As a stand-in for the processing done by the visual stream before the inferotemporal or perirhinal cortices, we use a pre-trained convolutional neural network (CNN), and extract the activity in its penultimate layer (before the final classification step). We use the ResNet18 network (He et al., 2015), although any CNN could, in principle, be used. This activity is a 512-dimensional vector, which, if used as the HebbFF input dimension *d*, would lead to the capacity *R*^max^ being prohibitively large for training purposes. To keep the performance in a reasonable range, we downsample to *d* = 50, either by partial sampling (Fig 8a) or by introducing an intermediate layer (Fig S6a).

**Figure 8.**
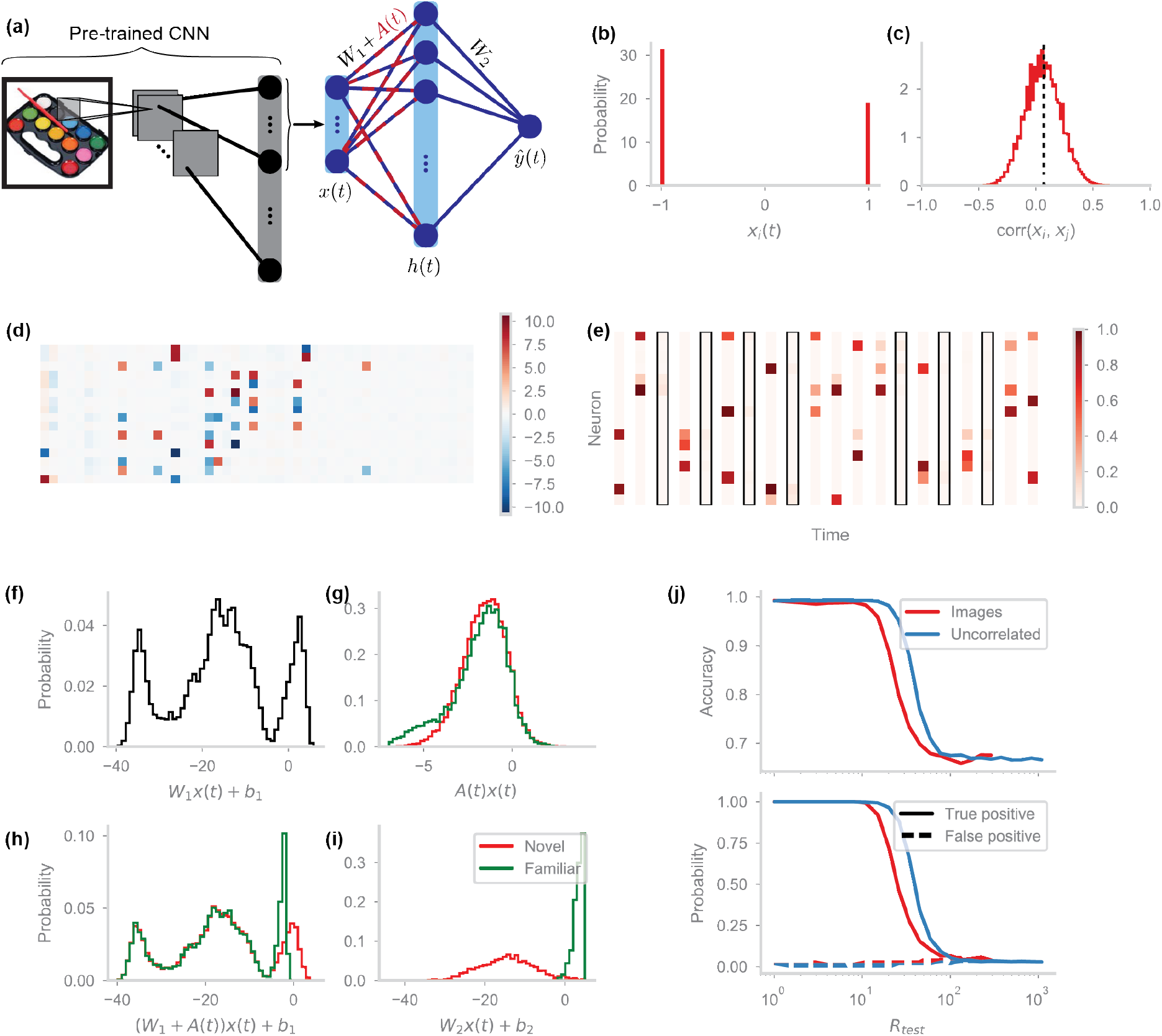
HebbFF performance on real-world images. (a) Network architecture for familiarity detection of real-world images. The activity of the penultimate layer of a convolutional neural network (ResNet18, pre-trained on ImageNet) is downsampled and passed to the HebbFF network (*d* = 50, *N* = 16) for familiarity detection. Only the HebbFF portion of this network is trained, via curriculum training. (b) Distribution of inputs ***x***(*t*) to HebbFF. After down-sampling by extracting the first 50 units of the CNN, the activity is centered at zero and binarized. (c) Histogram of the correlations between all pairs of input stimuli ***x***(*t*). On average (vertical dashed line) the correlation is slightly positive. (d-h) Same plots as Fig 4(f,c,i,j,m), respectively (*R*_train_ = *R*_test_ = 12). (j) Generalization performance, compared to a network of the same size trained on uncorrelated binary random vectors, is lower due to correlations in the input images.

As the first method of downsampling, we truncate the output of the CNN (Fig 8a). To keep the same input datatype as in previous sections, we also shift the inputs to zero mean and binarize them by taking the sign of each input component (Fig 8b). Unlike in previous sections, however, the inputs to HebbFF now have correlations that tend to be positive (Fig 8c). Nevertheless, this network has qualitatively similar features as the networks trained on uncorrelated vectors. The ***W***_1_ matrix has a similar structure (Fig 4f, 8d), the hidden layer activity is sparse (Fig 4c 8e), and the hidden unit input current distributions have similar shapes (Fig 4i,j,m 7b, 8f-i). Due to the added correlations, however, there is a decline in performance compared to a network of the same size trained and evaluated on uncorrelated binary random vectors (Fig 8j).

As another way to downsample, we add a trainable linear layer that transforms the CNN output to a 50-dimensional real-valued vector (Fig S6a). After training, the resulting inputs to HebbFF are no longer binary, but they are zero-mean (Fig S6b) and have zero-mean correlations Fig S6c). Interestingly, the network learns to generate this representation automatically to optimize familiarity detection over long intervals, which further supports storing uncorrelated stimuli. Although the ***W***_1_ matrix (Fig S6d) and the distribution of input currents from the fixed component of the synapses (Fig S6f) have a different structure compared to the original network, the operating principle remains the same: the ***W***_1_ matrix acts as a hash function to induce sparse activity in the hidden layer (Fig S6e) that is then suppressed for a familiar stimulus through the ***A***(*t*) matrix (Fig S6g-h). The network maintains its generalization performance across repeat intervals *R*, and across permutations of the sequence of images (Fig S6j). However, it does not generalize well to images it has not been trained on. It is possible that this difficulty is due to the relatively small number of images used during training and may be addressed by using a much larger dataset such as ImageNet (Deng et al., 2009).

## Discussion

Continual familiarity detection is a memory task that we perform every day, typically without being aware that we are doing it. We have used meta-learning to generate networks that solve this task using synaptic plasticity. This is distinct from memory storage in RNNs that maintain memory traces through persistent activity. Given the extraordinary capacity and robustness of recognition memory, the idea that biological networks use ongoing activity for this purpose appears untenable (Lundqvist et al., 2018; Masse et al., 2020). If a neuronal network is storing a stimulus by maintaining a particular firing rate pattern across its neurons, any other computation risks disrupting that memory trace. In contrast, storage through synaptic updates leaves the neuronal activity free to perform other computations unrelated to memory storage (Ba et al., 2016). In addition, we found that synaptic plasticity provides a better inductive bias than recurrence for familiarity detection. After optimization through meta-learning, the HebbFF network not only outperforms RNNs on the task, but also easily generalizes both in-distribution and out-of-distribution of the training data. Although RNNs are a common approach to tasks that require storage of the input history (Elman, 1991; Hochreiter and Schmidhuber, 1997; Mante et al., 2013), this result highlights the importance of considering alternative architectures and storage mechanisms, such as those that rely on synaptic, rather than neuronal, dynamics.

We found that anti-Hebbian plasticity, in which neuronal co-activation causes synaptic depression, is a better storage mechanism for familiarity detection than Hebbian plasticity. An anti-Hebbian rule generalizes better, has a larger capacity, and is discovered by meta-learning more frequently and reliably. Although this result is consistent with previous work (Bogacz and Brown, 2003), the underlying reasons are different. Bogacz and Brown showed that in a non-continual version of the familiarity detection task, an anti-Hebbian plasticity rule leads to a larger storage capacity, although this advantage only held in the case of correlated inputs. In their case, the anti-Hebbian rule automatically suppresses common input features, effectively storing only the uncorrelated components, leading to an increased capacity. In contrast, anti-Hebbian HebbFF shows an advantage even for uncorrelated inputs in the continual task. This is due to an effective decrease in the number of plasticity events – a synaptic update is weak for a familiar stimulus because the postsynaptic activity is low, leading to smaller updates that are less disruptive to stored memories.

In addition to reproducing prior experimental results such as repetition suppression, the HebbFF model allows us to make a novel experimental prediction. Although is obvious that the true positive rate (probability of correctly identifying a repeated stimulus as “familiar“) should decrease with longer delay intervals *R*, we also observe that the false positive rate slightly increases. Because anti-Hebbian plasticity causes repetition suppression, false negative responses arising from spurious activity in the hidden layer cause additional plasticity, depressing a subset of synapses. Subsequently, a novel stimulus is more likely to silence the hidden layer units, signaling familiarity and resulting in a false positive response. Prior experimental paradigms have not measured this effect because each trial had familiar stimuli interleaved at various delay intervals. As a result, novel stimuli could not be separated scored depending on the difficulty of the dataset, and false positive probability was reported in aggregate. If biological networks implement familiarity detection through an anti-Hebbian plasticity mechanism, we expect an increase in the false positive rate to coincide with a decrease in the true positive rate.

There are experimental results that the HebbFF model does not capture. For example, data from human subjects shows a very slow decrease in performance as a function of *R* that begins at relatively small value (Brady et al., 2008). In contrast, HebbFF has near-perfect performance for all *R* < *R*^max^, and then performance drops off quickly. However, it is likely that errors in the experiments do not reflect limitations on recognition memory but rather are due to factors such as fatigue and lack of attention that were not included in the model.

Finally, our work demonstrates the utility of meta-learning as a tool for neuroscience discovery. We used meta-learning to optimize a network architecture and plasticity rule that solves the continual familiarity detection task, contrasted it with an alternative sub-optimal solution, and subsequently used analytic methods to understand its mechanism. A similar approach can be used for other networks, plasticity rules, datasets, and tasks.

## Acknowledgement

We thank Stefano Fusi, Ken Miller, Dmitriy Aronov, James Murray, Marcus Benna, SueYeon Chung, Juri Minxha, Taiga Abe, and Denis Turcu for helpful discussions. Research supported by NSF NeuroNex Award DBI-1707398, the Gatsby Charitable Foundation, and the Simons Collaboration for the Global Brain. G.R.Y. was additionally supported by the Simons Foundation. We acknowledge computing resources from Columbia University’s Shared Research Computing Facility project, which is supported by NIH Research Facility Improvement Grant 1G20RR030893-01, and associated funds from the New York State Empire State Development, Division of Science Technology and Innovation (NYSTAR) Contract C090171, both awarded April 15, 2010.

## Online methods

### HebbFF and RNN training

To set the fixed HebbFF parameters ***W***_1_, ***b***_1_, ***W***_**2**_, ***b***_**2**_, *λ*, *η*, as well as the RNN weight and bias matrices, we use the PyTorch implementation of the Adam optimizer with the suggested default hyperparameters (Kingma and Ba, 2017). For a single trial, we use a dataset containing *T* stimuli, with familiar ones appearing at a repeat interval *R*. We present stimuli to the network sequentially, and compute the binary cross-entropy loss

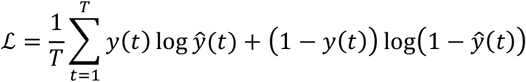

Since this is a dynamic task (the state of the network at time *t* + 1 depends on the state at time *t*), backpropagation through time is used to compute the gradient of the loss with respect to the parameters.

For each trial, we either use the same pre-generated length-*T* dataset, or we generate a new length-*T* dataset using the same repeat interval *R*. We refer to the latter case as the “infinite data” training regime since the sample space is much larger than the network would explore during training. Note that in the infinite data regime, we do not consider a validation dataset, since the training set is new every time and the training accuracy is therefore the same as the validation accuracy. In both cases, one trial corresponds to one step of gradient descent. To train the HebbFF network, the plastic matrix ***A***(*t*) is reset to a matrix of zeros at the start of each trial. Similarly, when training the RNN, hidden unit activity is reset to zero. In practice, the plastic matrix of HebbFF reaches its steady state distribution quickly and the transient does not contribute significantly to the gradient, so any reasonable initialization can be used.

To train the HebbFF network on the augmented familiarity detection/object classification task, we simply sum the cross-entropy losses from the classifier and familiarity output units:

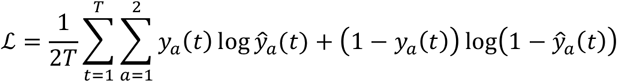

For every trial, we draw a new dataset from the pre-generated pool of stimuli. The class of each stimulus remains the same across datasets, but the ordering and repeats are chosen randomly each time. Although the network will have seen all of the stimuli during training in order to learn their classes, we can test generalization performance on the familiarity subtask by varying *R* and generating previously unseen permutations of the stimuli.

The PyTorch implementation of each of these can be found at https://github.com/dtyulman/hebbff.

### Bogacz-Brown (Bogacz and Brown, 2003) model implementation

To validate it on the (non-continual) two-alternative-forced-choice (2AFC) familiarity detection task, we implement the anti-Hebbian model as described by Bogacz and Brown (Bogacz and Brown, 2003), with the exception that the distribution of weights in the plastic weight matrix must be normalized such that its variance is equal to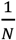,rather than unit variance as stated in the paper. In the encoding phase, the network is presented a sequence of *P* random patterns. In the testing phase, it is shown the original *P* patterns, as well as *P* novel ones. Critically, there are no plastic updates in the testing phase. A stimulus is reported as “familiar” if the output unit activity is below the mean across all 2*P* test patterns and “novel” otherwise. We see that this model performs well on the 2AFC task with a range of plasticity rates η (Fig S2a), so we arbitrarily choose η = 0.7 to test its performance on the continual task.

The continual task, unlike the 2AFC task, does not have an equal proportion of novel and familiar stimuli since we ensure that a stimulus is repeated at most once. So, we set the readout threshold such that an item is considered novel if it is in the *f*^th^ quantile of output unit activity for that trial, where *f* is the fraction of novel stimuli in the trial. This ensures that the fraction of stimuli reported as “novel” is equal to the true fraction of novel stimuli. In the case of equal proportions of novel and familiar stimuli, this reduces to the threshold being equal to the mean of the output unit activity for that trial.

Finally, note that unlike in the 2AFC task (Fig S2a), the performance of this model does not go to chance levels for large dataset sizes *T* in the continual task (Fig 3d). Rather, the true positive rate goes to zero and the false positive rate is ≈ 0.5, so accuracy is ≈ 0.33. The reason for this difference is that the second presentation of a stimulus in the continual task causes an additional plasticity event, unlike the 2AFC task where the test phase is offline. As a result, for datasets much larger than the network capacity *T* ≫ *P*\*, the output unit activity for familiar stimuli becomes larger than the activity for novel stimuli (Fig S2b).

### Training FLD and SCC decoders

To construct the Fisher linear discriminant (FLD) and spike count classifier (SCC) decoders, we first generate a dataset of length *T* = 1000. To better match the experimental dataset (Meyer and Rust, 2018), we use multiple values of *R* in this single stream. For each familiar stimulus, the value of *R* is drawn uniformly at random from 34 unique values, log-spaced from 1 to 100 (in practice, the results are qualitatively the same regardless of the number of items, the range, or whether the spacing is linear or logarithmic). We evaluate the trained network on this dataset and use the firing rates of the hidden layer to perform analyses analogous to those reported in (Meyer and Rust, 2018).

We compute the readout weight and bias terms for the FLD decoder as

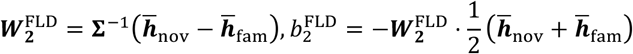

where 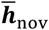 and 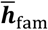 are the average firing rates of the hidden layer for novel and familiar stimuli, respectively, and the mean covariance matrix is calculated as

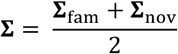

where Σ_fam_ and Σ_nov_ are the covariance matrices of the firing rates of the hidden layer for familiar and novel stimuli, respectively. The SCC decoder is a simple weighted average

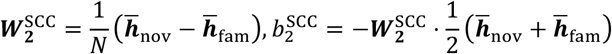

To get the ranking of the units for both decoders, we sort their readout weights and consider the most negative weights as the highest ranked. Note that for both decoders, the sign of the weights is flipped compared to (Meyer and Rust, 2018), and high-ranked units have the most negative weights rather than positive. This is due to the fact that we ask the network to label familiar stimuli as *y*(*t*) = 1, whereas (Meyer and Rust, 2018) readout a familiar stimulus as *y*(*t*) = 0. The two cases are symmetric and this does not change the results.

### Idealized model analytic capacity derivation

For notational simplicity, we only consider the nonzero submatrices of ***W***_1_ and ***A***(*t*), each of which acts on its corresponding subset of the input vector ***x***(*t*). Thus, equivalently, input layer of the idealized network is a *d*-dimensional vector split into two parts ***x***(*t*) = [***x***_W_(*t*),***x***_***A***_(*t*)], of dimension *n* and *D* respectively (*d* = *n* + *D*). Thus, the firing rate of the hidden layer is given by

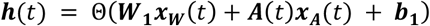

for an *N*×*n* matrix ***W***_1_, an *N*×*D* matrix ***A***(*t*), and an *N*×1 vector ***b***_1_. In other words, the firing rate of the *i*^th^hidden unit is

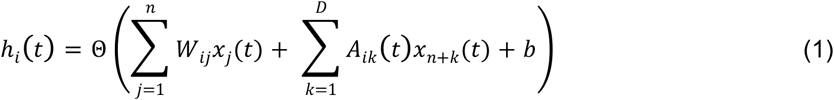

for *i* = 1,…,*N*, where Θ(·) is the Heaviside step function, i.e. Θ(Z) = 0 for Z < 0 and 1 for Z ≥ 0. We fix the value of *b* to be the same for all *i*. As before, the elements of ***x***(*t*) are +1 or −1 with equal probability. We would like to specify the network parameters such that exactly one hidden neuron is active for a novel stimulus and none for familiar, which will serve as the familiarity readout mechanism.

The *N*×*n* matrix ***W***_1_ is designed such that the vector ***W***_1_***x***_***W***_(*t*) has exactly one maximal entry given any such ***x***(*t*). Importantly, this matrix must act like a hash function such that different values of ***x***_***W***_(*t*) result in different entries of ***W***_1_***x***_***W***_(*t*) attaining the maximum value. One such ***W***_1_ is one whose rows enumerate all of the binary length-*n* strings consisting of entries +1 and −1. This sets the number of rows *N* to be equal to the total number of such strings, *N* = 2^*n*^. To set the overall scale of the input current (the term inside the nonlinearity), we scale this matrix by a factor *K*, to be determined later. For example, if *n* = 3,

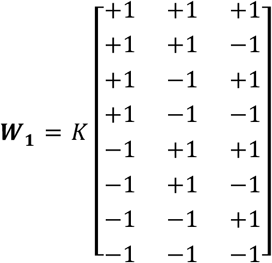

Thus, we have 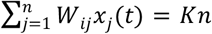 for exactly one value of *i*, specifically the row where *W*_*ij*_ = *x*_*j*_(*t*) for all *j*. This is the unique maximal value and will correspond to a different row for each instance of ***x***_***W***_(*t*).Subsequently, 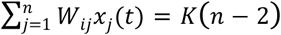 for *n* values of *i*, specifically those where *W*_*ij*_ ≠ *x*_*j*_(*t*) for exactly one *j*, and so on. Assuming that the vector ***A***(*t*)***x***_***A***_(*t*) is zero-mean with sufficiently small variance (this will be made rigorous shortly), we can now choose the scalar offset *b* such that exactly one element of ***h***(*t*) is equal to 1 and all others are zero.

The *N*×*D* matrix ***A***(*t*) is updated at every timestep by ***A***(*t* + 1) = λ***A***(*t*) − η***h*** *t****x***_***A***_(*t*)^T^, where the plasticity rate η is now restricted to be positive, corresponding to an anti-Hebbian learning rule. Considering one entry in this matrix and unrolling this recurrence, we find that

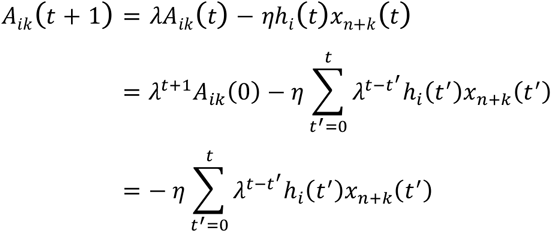

where the last equality holds if we assume that the network is in steady-state, so *t* is large, i.e. *t* → ∞, and therefore λ^*t*+1^*A*^*ik*^(0) → 0.

We can now consider the middle term of eq. 1, which we denote by ε_i_(*t*). We consider it as a random variable and compute its mean and variance. By definition, we have

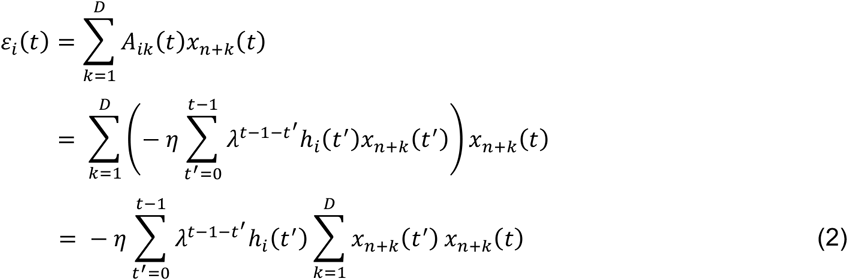

In the case where ***x***(*t*) is novel, *x*_*n+k*_(*t*′) and *x*_*n+k*_*t* are independent Bernoulli random variables that take on values ±1 with probability 1/2. Thus, X_*k*_(*t*′) = *x*_*n+k*_(*t*′)*x*_*n+k*_(*t*) is also a Bernoulli random variable with the same distribution, zero mean and unit variance, so

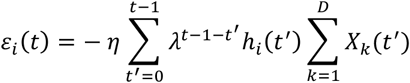

Since the entries of ***x***(*t*) are independent by definition, the X_*k*_(*t*′) are also independent across *k*, so summing over these indices, the variances add. Therefore,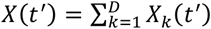 is a random variable with mean 0 and variance *D*, and

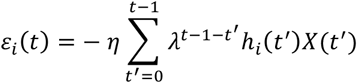

Next, we need the statistics of the term *h*_*i*_(*t*′). Since it is a function of the random variable ***x***(*t*), we also consider it as a random variable. Let *f*_eff_ denote the fraction of stimuli reported as “novel” by the network. Note that there are two ways for a network to report a stimulus as “novel” – by correctly identifying a novel stimulus (“true negative“), or incorrectly identifying a familiar one (“false negative“) – so if we let *f* denote the true fraction of novel input stimuli, we have

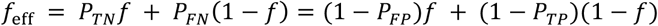

where *P*_*TN*_,*P*_*FN*_,*P*_*TP*_and *P*_*FP*_are the true negative, false negative, true positive, and false positive rates, respectively. Since by design there is exactly one hidden unit active for a novel stimulus, we have *h*_*i*_(*t*′) = 1 with probability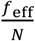, and *h*_*i*_(*t*′) = 0 with probability 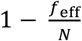. So, *h*_*i*_(*t*′) is a Bernoulli random variable with mean 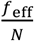 and variance 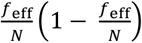. Now, we let *H*_*i*_(*t*′) = *h*_*i*_(*t*′)*X*(*t*′), so

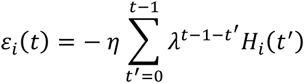

Although *h*_*i*_(*t*′) is, in principle, a function of ***x***(*t*′), we assume they are independent. Since *X*(*t*′) is zero-mean, the mean of *H*_*i*_(*t*′) is also zero. Using the identity 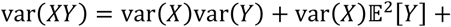 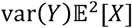, which holds for independent random variables *X* and *Y*, we have that the variance of *H*_*i*_(*t*′) is 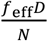. Finally, for convenience we can rewrite this as

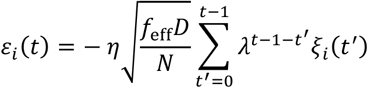

where ξ_*i*_(*t*′) is a zero-mean, unit-variance random variable. Furthermore, we now see that by the Central Limit Theorem ɛ_*i*_(*t*) is a Gaussian random variable since we are considering the steady-state performance at large *t*, so we can take *t* → ∞.

We can now compute the mean and variance of ɛ_*i*_(*t*). First, since *x*_*n+k*_(*t*) is zero-mean and independent of *A*^*ik*^(*t*),

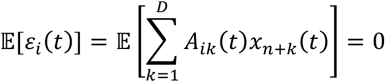

To compute the variance,

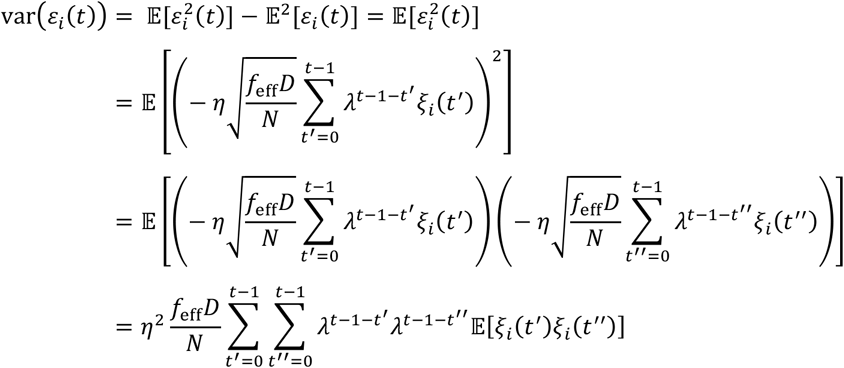

In general, we have 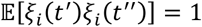 for *t*′ = *t*′′ since ξ_*i*_(*t*′) is a zero-mean, unit-variance random variable. For *t*′ ≠ *t*′′, we again make a simplifying independence assumption. In principle, ξ_*i*_(*t*′′) is not independent of ξ_*i*_(*t*′) since *h*_*i*_(*t*′′) depends on *h*_*i*_(*t*′) for *t*′′ > *t*′ through the memory stored in the ***A***(*t*) matrix. This dependence, however, is sufficiently weak, so we let 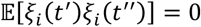 for *t*′ ≠ *t*′′. As a result, the double-sum collapses and we have

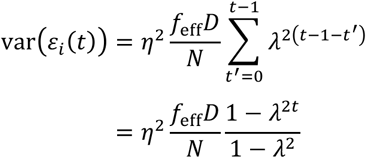

where the second equality comes from the standard geometric series. As before, since we are considering the steady-state with *t* → ∞, we have γ^2t^ → 0, so

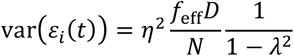

Thus, for a novel input ***x***(*t*) we can write

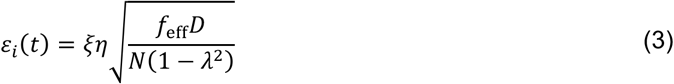

for all *i*, where ξ is a zero-mean, unit-variance Gaussian random variable, since ɛ_*i*_(*t*) is Gaussian.

For a familiar stimulus, where ***x***(*t*) = ***x***(*t* − *R*), clearly *x*_*n+k*_(*t*′) and *x*_*n+k*_(*t*) are no longer independent for *t*′ = *t* − *R*. Thus, we consider this term separately, rewriting the sum in eq. 2 as

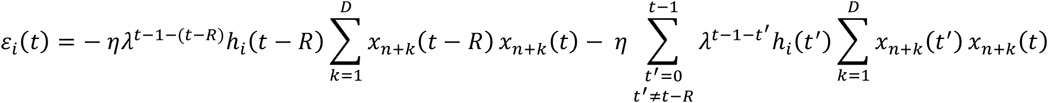

Assuming no errors, by design, *h*_*i*_(*t* − *R*) = 1 for exactly one neuron *i*, since the stimulus at time *t* − *R* was guaranteed to be novel (we enforce that a stimulus is repeated at most once in this task). We consider the statistics of ɛ_*i*_ *t* for this particular neuron. In the first term, the sum 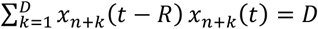 since by assumption *x*_*n+k*_(*t*) = *x*_*n+k*_(*t* − *R*) for all *k*. The second term has the same distribution as the one for a novel input since we have only removed one term from the sum and *t* is large. Thus, for a familiar stimulus we can write

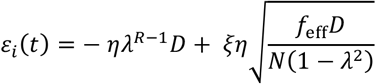

for exactly one value of *i*, where ξ is a zero-mean, unit-variance random variable as before. For all other values of *i*, eq. 3 holds.

Having established the statistics of the hidden layer input currents for a novel and a familiar stimulus, we can now write down the conditions for the model to work, use them to find the optimal values of the parameters and calculate the true positive and false positive probabilities, and compute the capacity – the largest value of *R* for which the error is below a predetermined threshold. First, to ensure that exactly one unit is active for a novel stimulus (true negative), since we are using a step function nonlinearity, we must have the largest input current take on a positive value,

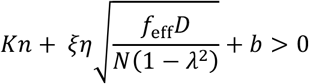

and second-largest to be below zero,

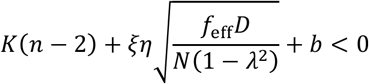

Second, to ensure there are no units active for a familiar stimulus (true positive),

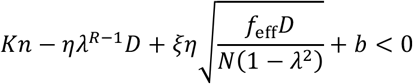

For sufficiently large *R*, i.e. if ηλ^R−1^*D* < 2*K*, the third of these conditions implies the second. Since we are interested in maximizing *R*, we only consider the first and third conditions. Furthermore, note that these conditions are overparameterized. If we divide all three equations by η (e.g. let 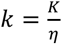, 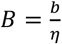) we can eliminate this free parameter. In other words, for any value of η we can scale *K* and *b* proportionally to satisfy the conditions, so for simplicity we choose η = 1. Similarly, the term *K*_*n*_ + *b* can be replaced by a single parameter since for any choice of *K* we can rescale *b* to keep this sum constant. To ensure that the condition ηλ^R−1^*D* < 2*K* holds for all *R*, we can choose *K* = *D*. For convenience, we also let *b* = β*D* and 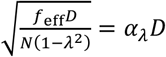, the subscript indicating explicit dependence on λ. Dividing both inequalities by *D*, the conditions simplify to

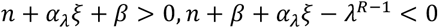

The accuracy, i.e. probability of a correct response, is given by *P*_*correct*_ = (1 − *f*)*P*_*TP*_ + *fP*_*TN*_. For convenience, we compute the false positive instead of the true negative rate, noting that *P*_*TN*_ = 1 − *P*_*TP*_. The false positive and true positive rates are given by

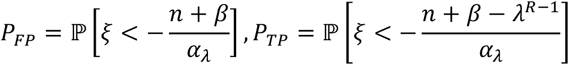

Since ξ is a standard Normal random variable, 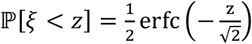, so

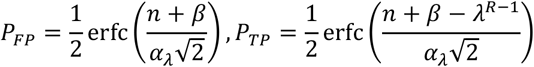

We would now like to set the optimal values of λ and β which maximize *R*, given a desired true positive and false positive probability 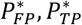. Note that fixing these probabilities also fixes 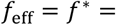 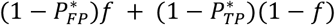. Rearranging the previous equations, we get

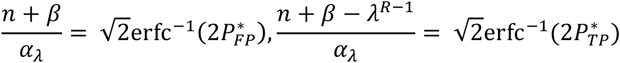

The first equality sets the value for β. To determine λ, we substitute β into the second equality to get

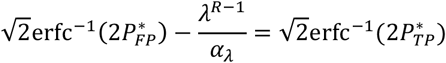

For notational convenience, let 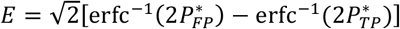. Using the definition of α_λ_ and *f*_eff_ = *f*^*^, we have 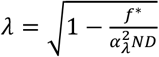. Rearranging, we have

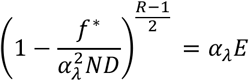

Assuming *N* and *D* are large (so λ is close to 1), we can use the first-order Taylor expansion exp(−Z)≈1 − Z for the term in parentheses (this will be necessary to get a closed-form expression for the optimal λ) and solve for *R*

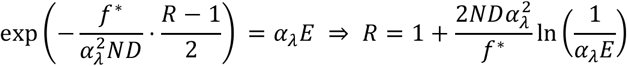

Setting 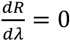 and solving for λ gives the optimum

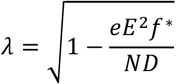

Thus, the capacity of the optimized network is

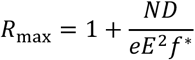

where *f*^*^ and *E* are constants that depend on 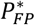 and 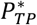 (*f*^*^ also depends on the true fraction of novel stimuli *f*). For instance, if we impose that 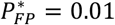 and 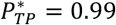, with our value of 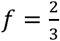, we get

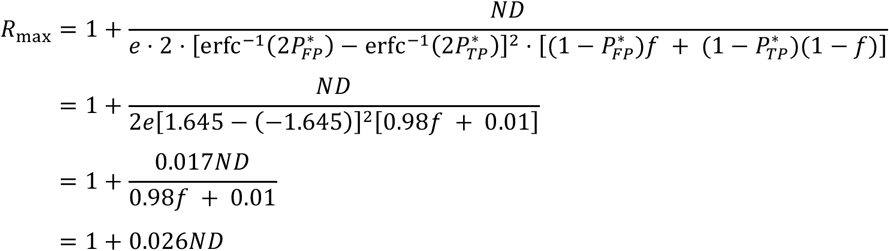

It is clear that that the capacity scales in proportion to the number of plastic synapses in the network. Furthermore, since *d* = *n* + *D*, i.e. *D* = *d* − log_2_(*N*), the capacity scales in proportion to the total number of synapses *d*, as long as *D* ≫ *n*.

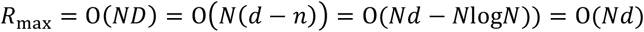

Finally, note that the equations for *P*_*FP*_and *P*_*TP*_are a function of *f*_eff_ due to the α_λ_ parameter, and therefore recursively depend on *P*_*FP*_and *P*_*TP*_. We cannot compute the closed-form solution for these, but we can approximate the values with arbitrary accuracy by iterating through this recurrence until convergence to the fixed point. As the initial value for the recurrence, we use *P*_*FP*_and *P*_*TP*_computed using *f*_eff_ = *f*, i.e. assuming no errors.

## Supplementary figures

**Figure S1.**
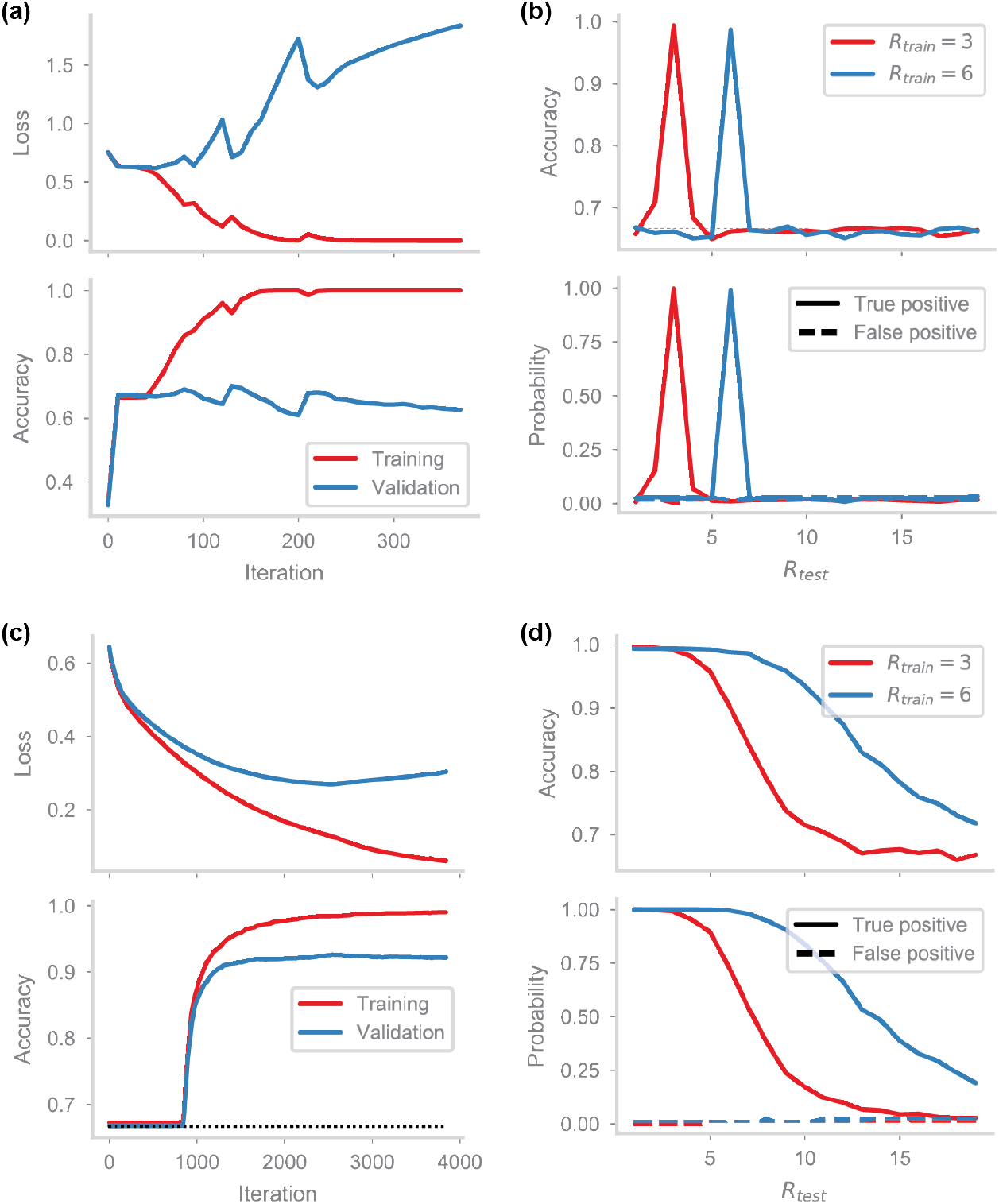
HebbFF and RNN comparison, matching total number of dynamic variables. (a,b) RNN performance as in Fig 2a-b, with *d* = 25 and *N* = 625. (c,d) HebbFF performance as Fig 3a-b, with *d* = 25 and *N* = 25. The number of plastic synapses in the HebbFF network *N* * *d* = 625 is the same as the number of recurrent units in the RNN, matching the number of dynamic variables between the networks rather than the number of neurons. HebbFF still shows better generalization in both training scenarios.

**Figure S2.**
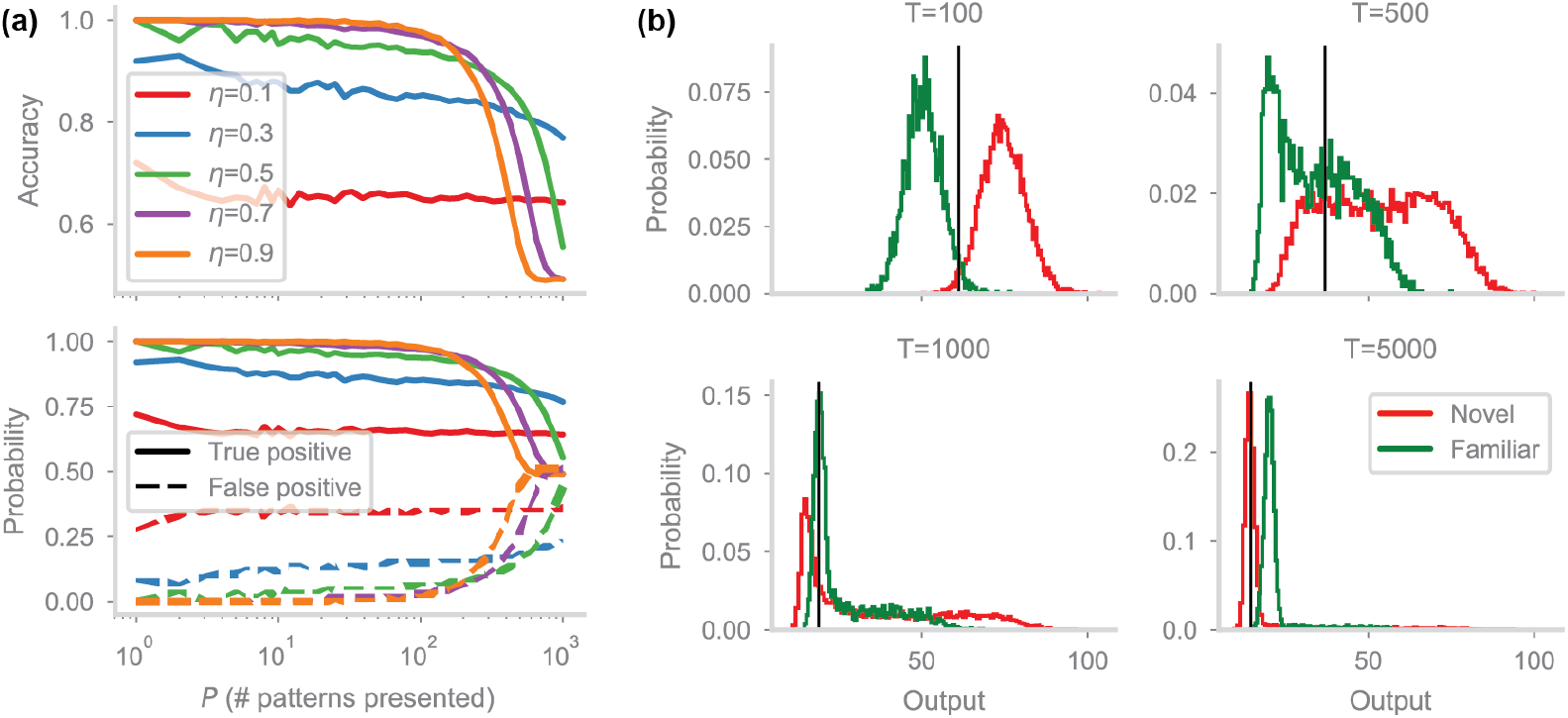
Validation of network from Bogacz and Brown (2003). (a) Performance of the anti-Hebbian network from (Bogacz and Brown, 2003) (*d* = *N* = 100) on the non-continual familiarity detection task. The network is shown *P* uncorrelated randomly generated patterns, and tested on all the presented patterns as well as *P* novel ones. If the network’s readout falls below a threshold, the corresponding input is classified as “familiar,” otherwise “novel.” The threshold is set such that that exactly half of the test inputs are classified as “familiar.” The plot shows the network’s performance for a range of learning rates η as a function of the number of patterns presented. (b) Performance of the same network (η = 0.7) on the continual task. Plots show the probability distribution of the network’s readout for novel (red) and familiar (green) stimuli. Threshold for familiarity (vertical black line) is set such that the top *f* of outputs are classified as novel (fraction *f* equal to proportion of novel stimuli in dataset), analogous to that in (Bogacz and Brown, 2003). Network performance declines as the distributions get closer together for increasing dataset size *T*. For very large dataset sizes, e.g. *T* = 5000, the readout for familiar stimuli is actually higher than that for novel, which causes overall performance (Fig. 3d) to fall below chance.

**Figure S3.**
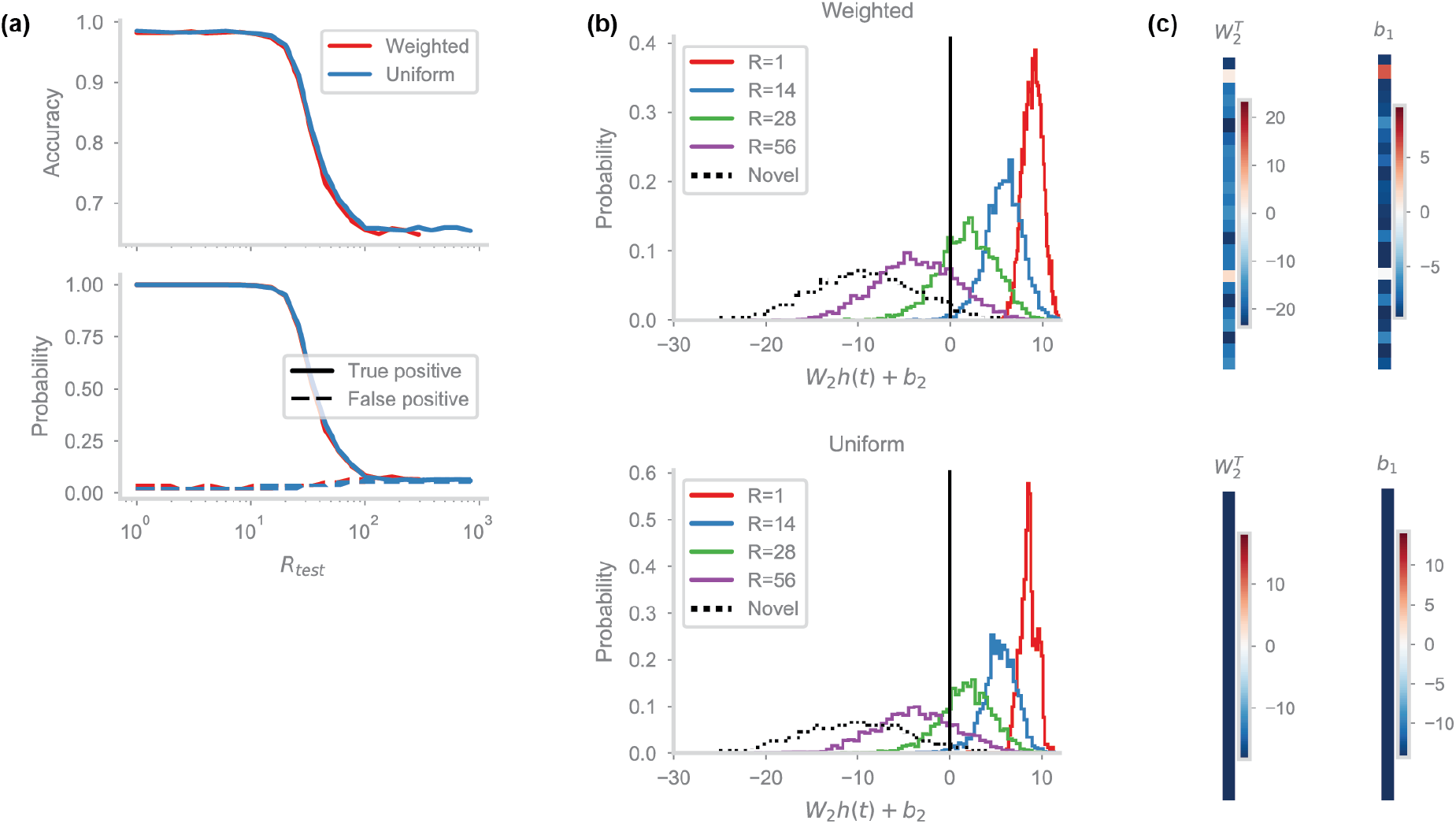
Comparison of uniform and trained readout. (a) Generalization performance of a trained HebbFF network (*d* = *N* = 25) with a fully trainable readout matrix, i.e. a weighted average of the hidden layer activity (red) and a version where all the entries of the readout matrix are constrained to be equal, i.e. a scaled average of the hidden layer (blue). (b) Distributions of the output unit activity (prior to applying the nonlinearity) for a trained network, for several test values of *R*. The output distributions, as well as performance, are almost identical for both versions of the readout. (c) Examples of the ***W***_**2**_ readout matrix, as well as the bias ***b***_1_ for the weighted (top) and uniform (bottom) readouts. Values for both are negative, indicating a qualitatively similar readout mechanism. Note that the weight matrices are plotted transposed for visualization.

**Figure S4.**
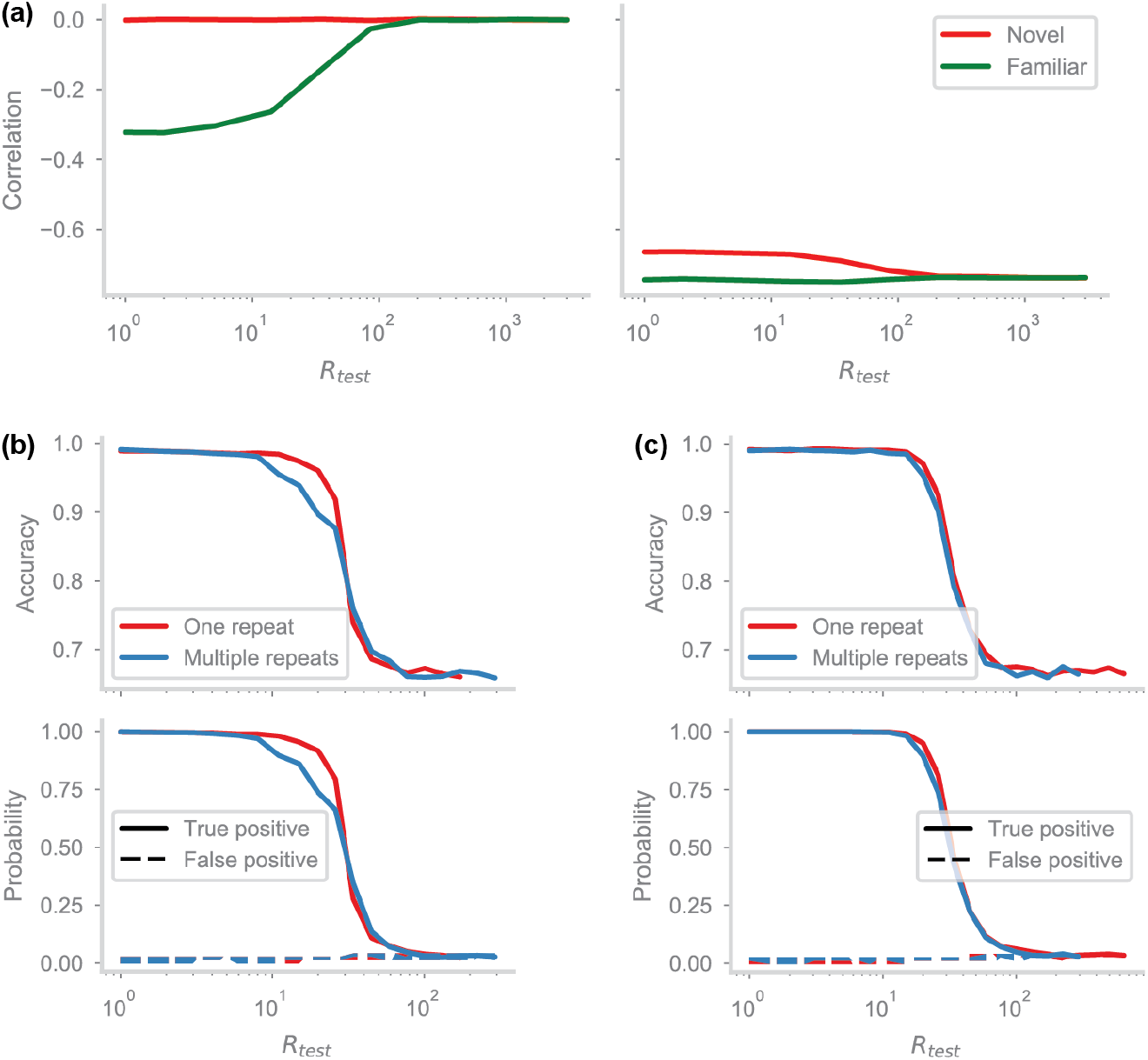
Idealized and HebbFF model differences. (a) Correlation between the hidden layer input currents due to static and plastic synapses for the idealized model (left) and HebbFF (right) for novel (red) and familiar (green) stimuli as a function of delay interval (*d* = 25, *N* = 32). In the idealized model, due to the split static and plastic synapses, the currents are uncorrelated for novel stimuli and familiar stimuli at long delay intervals. At short intervals, the two input currents are anti-correlated, which enables repetition suppression. In the HebbFF model, there is anti-correlation in both cases, although there is still less anti-correlation for novel stimuli. (b) Performance of the idealized network on the continual familiarity detection task with familiar stimuli repeated either exactly once, as used throughout this work (red), or multiple times (blue). Since there is exactly zero hidden unit activity for a familiar stimulus it does not get reinforced in memory, and less likely to be recognized on its second and subsequent repetitions. (c) The HebbFF network, trained to maximum capacity on the single-repeat task does not suffer any loss in performance due to multiple repeats.

**Figure S5.**
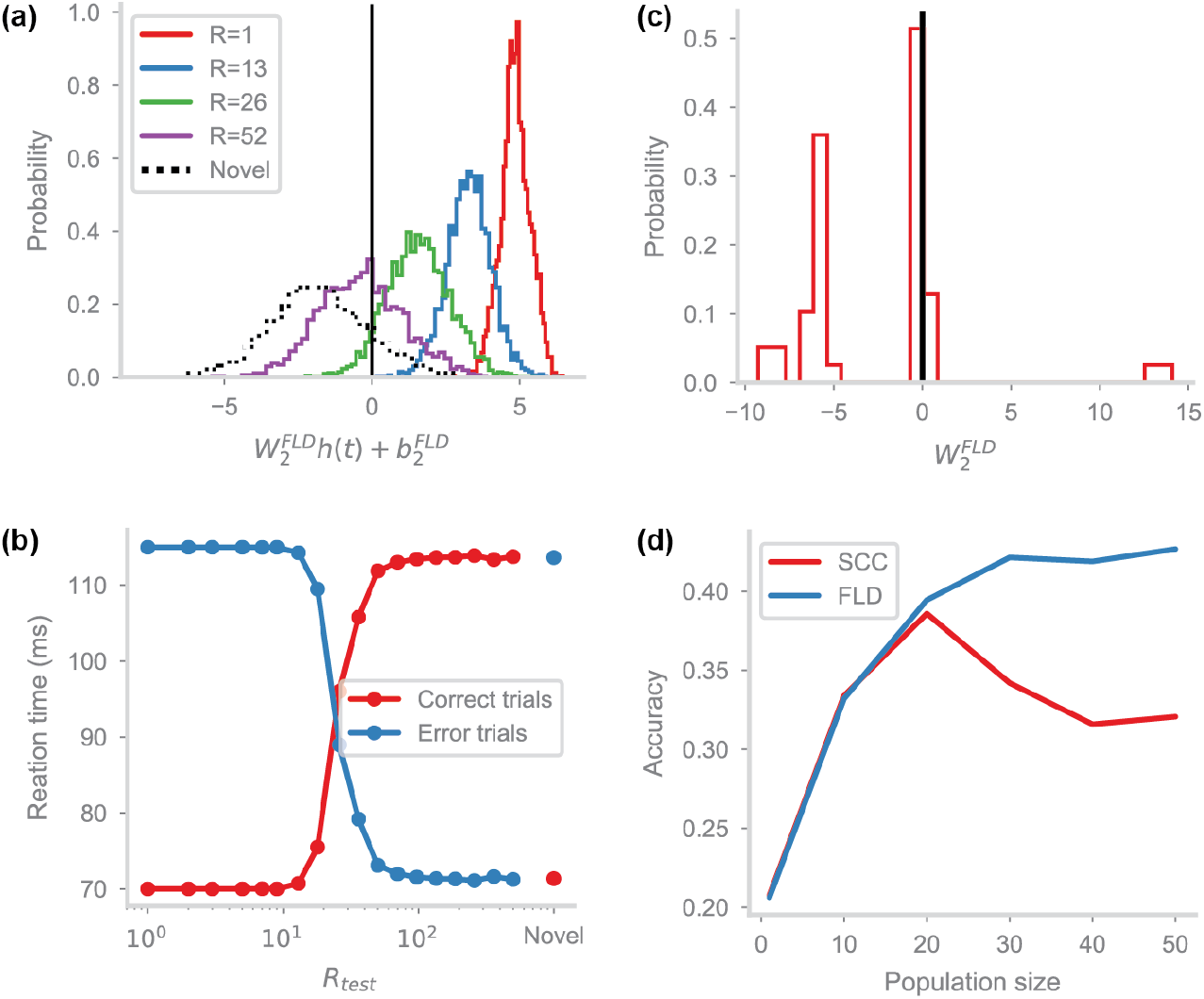
Behavior on augmented task. Panels correspond to the right-hand side plots of Fig 7(a-d), but for the classifier-augmented HebbFF network (*d* = 25, *N* = 50) performing binary classification and familiarity detection simultaneously.

**Figure S6.**
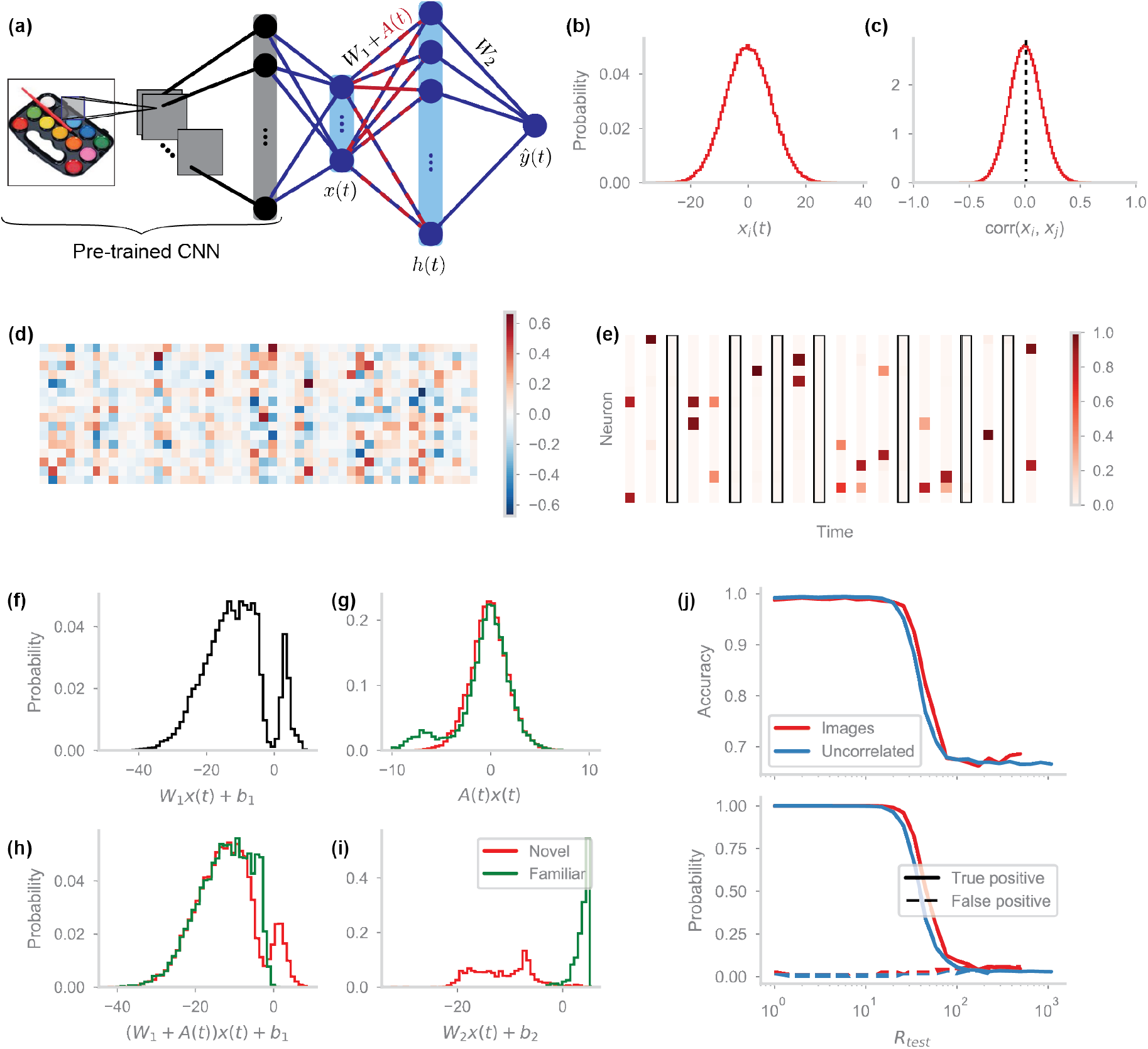
Performance on real-world images. Subplots as in Fig 8, but using a trained fully-connected linear layer to transform the activations of the CNN’s penultimate layer into the inputs of HebbFF (*R*_train_ = *R*_test_ = 19).

